# The Batten disease gene product CLN6 is a glycerophosphoglycerol epimerase that establishes *S,S*-BMP stereochemistry

**DOI:** 10.64898/2026.10.07.757477

**Authors:** Zhijun Zhu, Jian Xiong, Ibrahim Maaz, Matthew J. Matrongolo, Ali Goochani, Chetan Chilkunda, Xuejun Peng, Anish Alur, Christian Porsen, Angela Schulz, Susan L. Cotman, Natalia Gomez-Ospina, Monther Abu-Remaileh

## Abstract

Bis(monoacylglycero)phosphate (BMP), a lysosomal phospholipid essential for lipid catabolism and cholesterol clearance, has a noncanonical *S,S* glycerol configuration found in no other mammalian phospholipid, a feature necessary for its stability in the lysosome. How cells invert glycerol stereochemistry to build *S,S*-BMP has remained unexplained since this configuration was discovered five decades ago. Here, we show that the endoplasmic reticulum-resident Batten disease protein CLN6 is a Ca^2+^-dependent epimerase that isomerizes glycerophosphoglycerol (GPG) from *R,S*-GPG to *S,S*-GPG. CLN6-deficient cells, mouse tissues, and patient samples predominantly contain *R,S*-GPG, *S,R*-lysophosphatidylglycerol (LPG), and *R,S*-BMP instead of their *S,S* counterparts, with a net depletion of BMP. Purified CLN6 catalyzes GPG epimerization in vitro, an activity abolished by a pathogenic substitution in its substrate-binding pocket. Consistently, exogenous *S,S*-GPG restores BMP synthesis and relieves lipid storage in *CLN6*-null cells. Our work establishes stereochemical commitment upstream of acylation, placing CLN6, CLN8, and CLN5 in sequence in BMP biosynthesis and unifying the lipid defect across these neurodegenerative disorders.

## Introduction

Lysosomes catabolize lipids and recycle their content (*1*). Disruption of lipid recycling in the lysosome is implicated in a myriad of monogenic lysosomal storage disorders (LSD) in which genes encoding lipid hydrolases, their cofactors or lipid trafficking and transporting proteins are mutated (*2*). Emerging genetic and biochemical evidence further implicates lysosomal lipid metabolism in common neurodegenerative diseases including frontotemporal dementia (FTD), Parkinson’s, and Alzheimer’s diseases among others (*3–5*). Key to lipid catabolism in the lysosome is a unique lipid called bis(monoacylglycero)phosphate (BMP) (*6*). BMP lipids represent seventy percent of the phospholipid mass of intralysosomal vesicles (ILVs) (*7*), where they associate with lipid hydrolases and cofactors to facilitate proper lipid degradation (*8–12*) and sorting including cholesterol egress from the lysosome (*13–16*). It is because of these potential functions, boosting BMP levels is thought to be a promising therapy in lysosome-associated diseases where lipid clearance is perturbed (*6*).

BMP was discovered in 1967 by Body & Gray (*17*). BMP has unique stereochemistry; while most glycerophospholipids are assembled on an *sn*-glycero-3-phosphate backbone whose central glycerol carbon has the *R* configuration, BMP contains two glycerol moieties linked to a shared phosphate group through their respective *sn*-1 positions, giving rise to its characteristic noncanonical *S,S* configuration. This distinctive stereochemistry was recognized more than five decades ago, yet its biosynthetic origin has remained unresolved (*18, 19*). Earlier studies demonstrated glycerol reorientation during the conversion of phosphatidylglycerol (PG) to BMP, implicating a stereochemical transformation in the pathway (*20*). However, neither the enzyme responsible nor the intermediate at which this transformation occurs was established.

The Batten disease protein CLN5 was recently established as the lysosomal BMP synthase that catalyzes a reaction between two lysophosphatidylglycerol (LPG) molecules (*21*). Loss of CLN5 depletes BMP and causes accumulation of its LPG precursor, linking perturbation of BMP synthesis to neuronal ceroid lipofuscinosis (NCL). Subsequent work implicated the endoplasmic reticulum (ER) membrane protein CLN8, encoded by another NCL gene, in BMP biosynthesis by mediating the selective acylation of the *S*-glycerol carbon of glycerophosphoglycerol (GPG) to generate LPG, the direct substrate of CLN5 (*22–24*).

Like *CLN5* and *CLN8*, mutations in *CLN6* were identified as a cause of NCL more than two decades ago (*25*), but its function has remained undefined. CLN6 is an integral endoplasmic reticulum (ER) membrane protein with seven transmembrane domains. Interestingly, CLN6 has been shown to form a complex with CLN8 and is proposed to be critical for lysosomal function (*26, 27*), suggesting a potential role of CLN6 in BMP biogenesis.

Here we show that CLN6 loss leads to BMP depletion in human cell lines, induced pluripotent stem cells (iPSCs)-derived neurons, mouse, and patient samples. Mechanistically, we show that CLN6 is a Ca^2+^-dependent GPG epimerase that catalyzes the stereo-inversion of *R,S*-GPG to *S,S*-GPG. The catalytically dead pathogenic mutant of CLN6 abolishes activity, and exogenous *S,S*-GPG supplementation rescues BMP levels in *CLN6*-null cells. Our findings define the stereochemical transformation in *S,S*-BMP synthesis at the GPG stage and connect CLN6-dependent epimerization to the subsequent reactions mediated by CLN8 and CLN5. Together, these results explain how cells generate the distinctive stereochemistry of BMP and reveal a biochemical link between CLN6 dysfunction and defective lysosomal lipid biosynthesis.

## Results

### CLN6 loss reduces BMP levels

CLN3, CLN5 and CLN8 loss cause depletion of BMP lipids (*21, 23, 24, 28, 29*). We hypothesized that CLN6, reported to be in complex with CLN8 in the ER (*27*), is also involved in lysosomal lipid metabolism. To this end, we generated *CLN6*-knockout HEK293T cells using CRISPR-Cas9 (**fig. S1A**). Untargeted lipidomics revealed the reduction of BMP lipids as the major phenotype, accompanied by modest increase in cholesteryl esters (ChEs) and LPG, which is the BMP precursor (**Fig. 1A & Table S1**). Targeted analyses confirmed a broad reduction in BMP species, which was significantly reversed by re-expressing wild-type (WT) CLN6 but not the patient variants R62H and H127Y (**Fig. 1B & fig. S1B-D**). To further validate these results, we tested BMP levels in induced pluripotent stem cells (iPSCs) and their derivatives (**fig. S1E**). As expected, BMP levels were reduced in two *CLN6*-knockout iPSC clones (**fig. S1F**) and their differentiated neurons (iNeurons) (**Fig. 1C**). Whereas BMP levels were consistently reduced across these cell types, targeted analyses show that LPG levels increased modestly in *CLN6*-knockout HEK293T cells and iPSCs (**fig. S1, G and H**) but decreased in iNeurons (**fig. S1I**), indicating a dependence on cell type and media composition. Altogether, our results link CLN6 to BMP biology as in the case of several other NCL genes.

**Figure 1.**
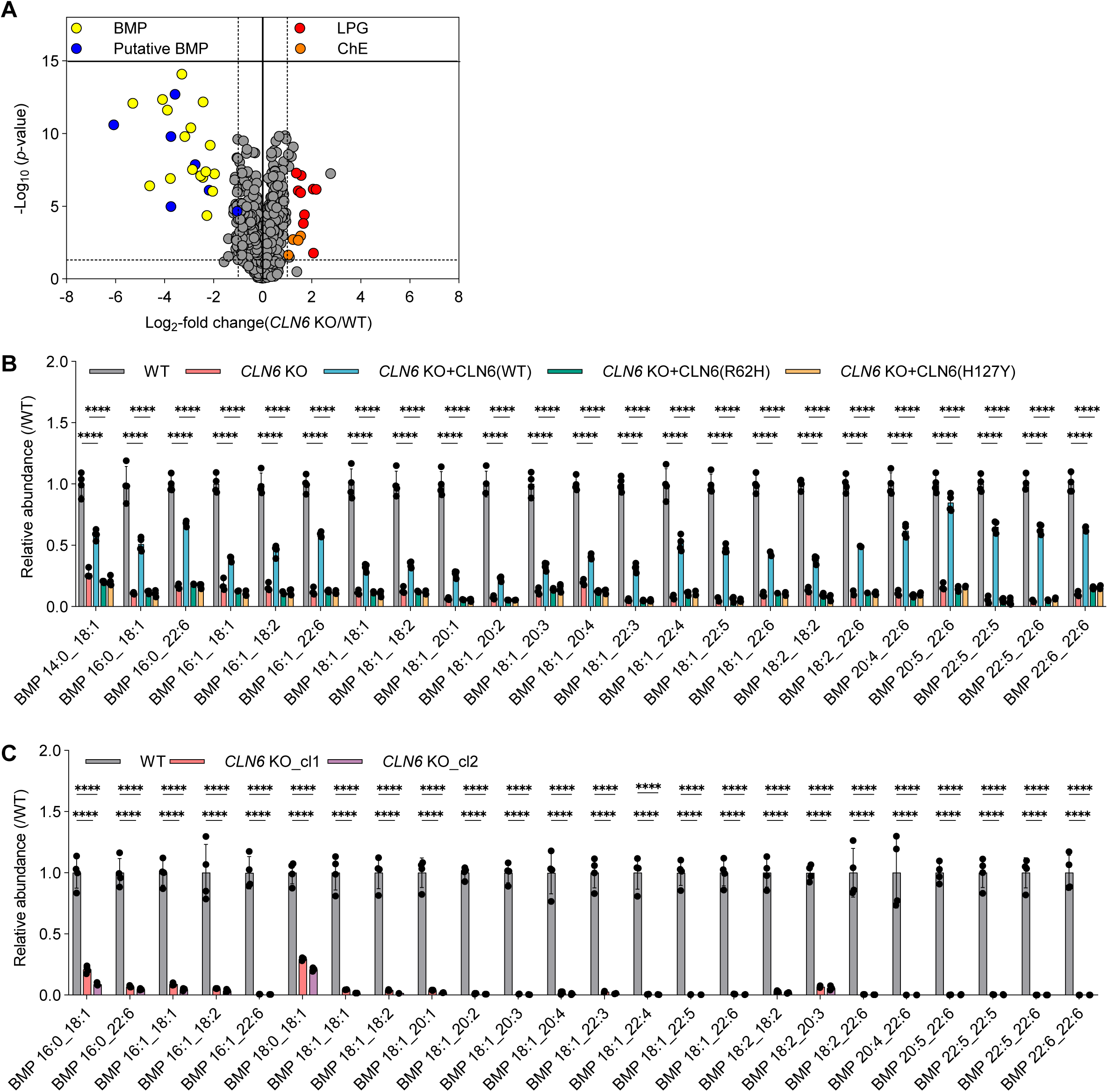
CLN6 loss reduces BMP levels. **(A)** Volcano plot of untargeted lipidomics data comparing *CLN6*-knockout (KO) and wild-type (WT) HEK293T cells. Each point represents a lipid feature, plotted by log_2_-transformed fold change in abundance (KO/WT) and −log_10_-transformed *p*-value. BMP, putative BMP, LPG, and cholesteryl esters (ChE) species are highlighted in yellow, blue, red, and orange, respectively; other features are shown in gray. Dashed lines indicate abundance ratios of 0.5 and 2, and *P* = 0.05. Putative BMP species were annotated on the basis of negative-ion MS/MS spectra and retention times but lacked MS/MS spectra of [M+NH] precursor ions. Each genotype measurement represents *n* = 6 biologically independent samples. *P*-values were calculated by one-way analysis of variance (ANOVA) with Tukey honestly significant difference (HSD) test and corrected by the Benjamini-Hochberg method. **(B)** Targeted lipidomics analysis of BMP species in WT HEK293T cells, *CLN6* KO cells, and KO cells re-expressing WT CLN6 or patient-derived variants R62H or H127Y. Data are presented as mean ± SD; *n* = 3 biologically independent samples for the *CLN6* KO group and *n* = 4 for each remaining group. **(C)** Targeted lipidomics analysis of BMP species in WT iNeurons and two independent *CLN6* KO derived from two independent iPSC clones (cl1 and cl2). Data presented as mean ± SD of *n* = 4 biologically independent samples. In **(B and C)**, the abundance of each BMP species was first normalized to endogenous POPC and then normalized to its mean abundance in the corresponding WT group. Points represent individual biological replicates. Statistical significance was assessed using two-way ANOVA; \*\*\*\**P* < 0.0001.

### CLN6 is required for *S,S* stereochemistry of BMP and LPG

The reduction, rather than complete loss, of BMP in the CLN6-deficient cells prompted us to investigate whether CLN6 loss alters its characteristic *S,S* stereochemistry, necessary for BMP resistance to phospholipase activity in the lysosome (*30*). Esterification of one of the two primary hydroxyl groups of glycerol with phosphate creates a chiral center at the central carbon. Under the stereospecific numbering (*sn*) convention, glycerol phosphate with phosphate at *sn*-3 has the *R* configuration, whereas its enantiomer with phosphate at *sn*-1 has the *S* configuration. The remaining hydroxyl groups occupy the *sn*-1 and *sn*-2 positions or the *sn*-3 and *sn*-2 positions, respectively (**Fig. 2A**).

**Figure 2.**
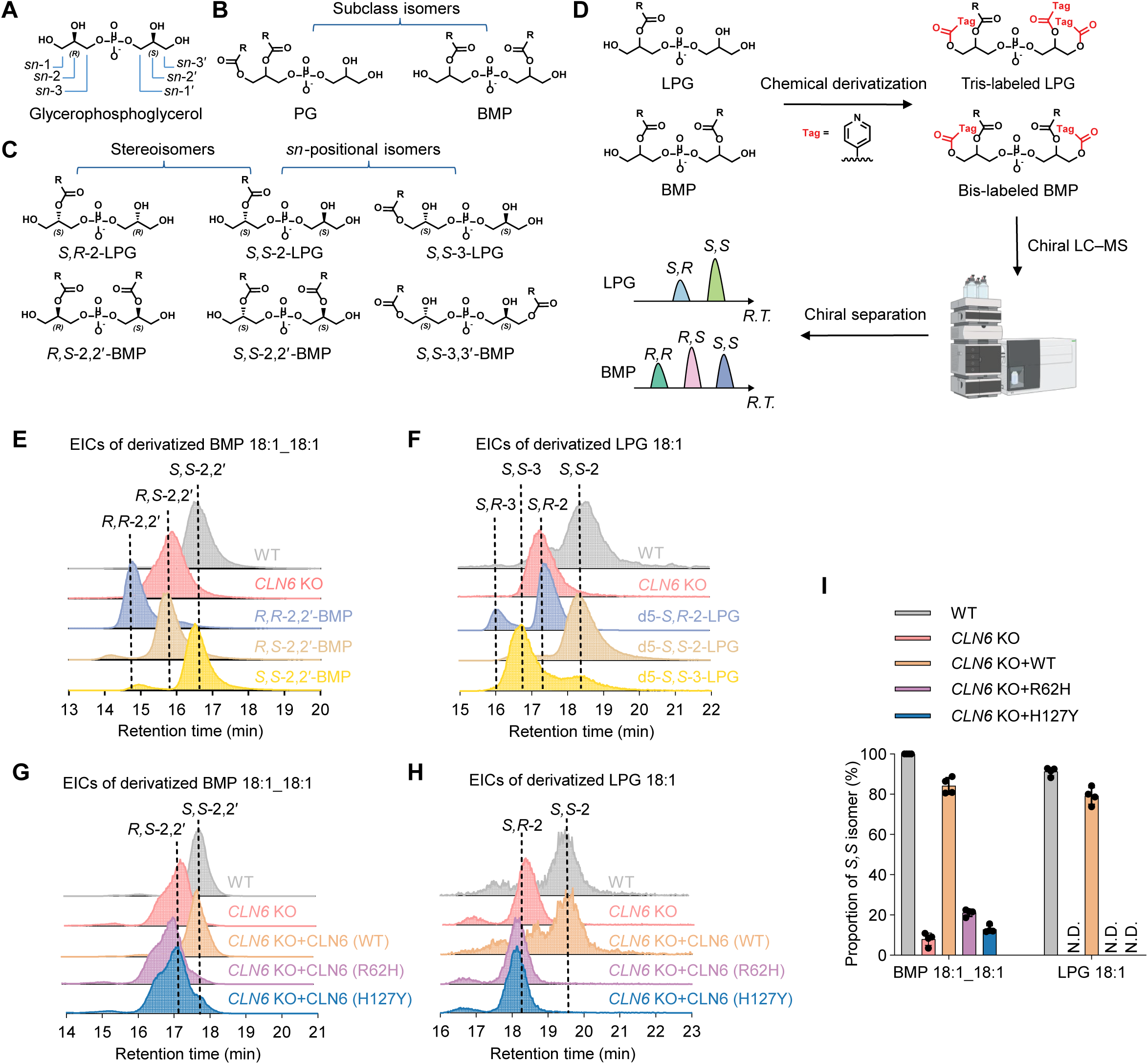
CLN6 is required to establish the *S,S* stereochemistry of BMP and LPG. (**A-C**) Chemical structures of glycerophosphoglycerol (GPG), LPG, PG, and BMP, illustrating glycerol stereochemistry and stereospecific numbering (*sn*) and representative stereoisomers and *sn*-positional isomers of LPG and BMP. **(D)** Workflow combining chemical derivatization of free hydroxyl groups with chiral LC–MS to resolve LPG and BMP stereoisomers and positional isomers. **(E and F)** Representative extracted ion chromatograms (EICs) of derivatized BMP 18:1_18:1 **(E)** and LPG 18:1 **(F)** from WT and *CLN6* KO HEK293T cells, shown alongside synthetic standards. LPG standards are d5-labeled. Dashed lines indicate the retention times of the indicated isomers. **(G and H)** Representative EICs of derivatized BMP 18:1_18:1 **(G)** and LPG 18:1 **(H)** from WT cells, *CLN6* KO cells, and KO cells re-expressing WT CLN6 or either patient-derived variant, R62H or H127Y. Dashed lines indicate the retention times of the indicated isomers. In **(E), (F), (G),** and **(H)**, each trace is independently normalized to its maximum peak intensity (100%). **(I)** Proportions of the *S,S* isomers of LPG 18:1 and BMP 18:1_18:1 in the cell groups shown in **(G and H)**. Data are presented as mean ± SD. *n* = 4 biologically independent samples per group. Panel D was created using BioRender (https://biorender.com).

Established LC–MS-based lipidomics workflows can reliably distinguish BMP and its structural isomer, phosphatidylglycerol (PG) (**Fig. 2B**) by their retention times and characteristic MS/MS fragmentation patterns (*31*). However, it remains challenging to resolve stereoisomers (*R* versus *S*) and *sn*-positional isomers (*sn*-1/*sn*-3 versus *sn*-2), particularly in biological samples (**Fig. 2C**). To this end, we developed a strategy combining chemical derivatization with chiral separation (**Fig. 2D** and see methods) to resolve the stereo– and regiochemistry of BMP 18:1_18:1 (**fig. S2A**) and LPG 18:1 (**fig. S2B**), relatively abundant species, validated by synthetic standards. By comparing retention times of endogenous BMP 18:1_18:1 to those of synthetic standards, we confirmed *S,S*-2,2 ′ –BMP as the predominant isomer in WT cells, which is lost in *CLN6* knockout cells, leaving only the *R,S* stereoisomer (**Fig. 2E**). The location of BMP in the lysosome while CLN6 is an ER-resident protein, rules out direct effect on BMP and suggests that the stereochemistry of LPG, which has been shown to be synthesized by CLN8 in the ER (*23, 24*), might be also impacted by CLN6 loss. Indeed, while *S,S*-2-LPG 18:1 is the predominant isomer in WT cells, *CLN6* knockout cells only have the *S,R* stereoisomer (**Fig. 2F**). Of importance, re-expressing WT CLN6 restored the *S,S* configuration of BMP and LPG, whereas neither patient-derived variant did (**Fig. 2G-I**). Altogether, these results demonstrate that CLN6 is required to establish the *S,S* stereochemistry of both LPG and BMP.

### CLN6 is required to generate *S,S*-GPG

Recent reports indicate that acylation of GPG by CLN8 to form LPG occurs on the glycerol moiety with the *S* configuration (*24*). We therefore hypothesized that CLN6 is required for the isomerization of *R,S*-GPG to its epimer *S,S*-GPG, which then incorporates in the BMP biosynthesis pathway (**Fig. 3A)**.

**Figure 3.**
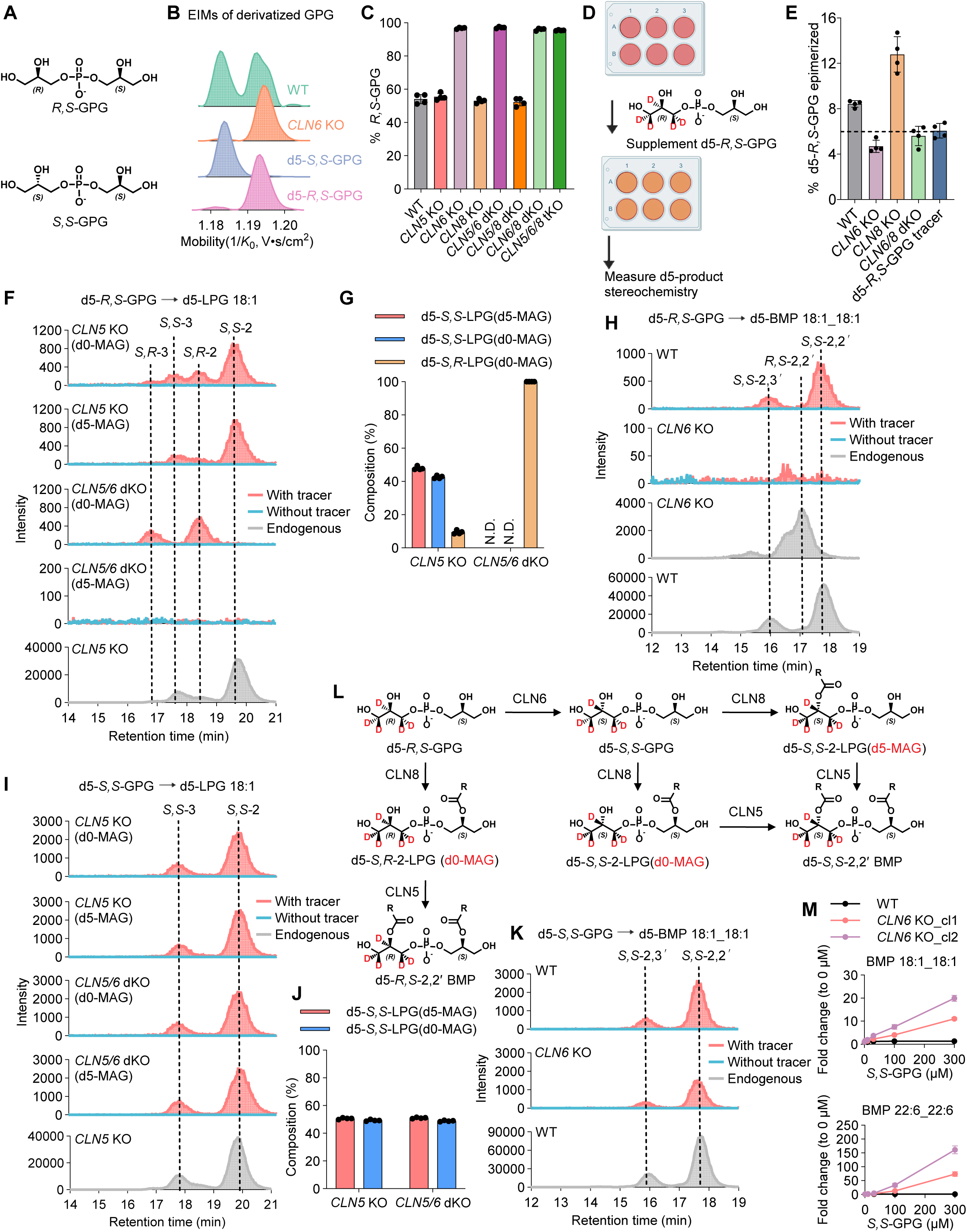
CLN6 is required to generate *S,S*-GPG. **(A)** Chemical structures of *R,S*-GPG and *S,S*-GPG. **(B)** Representative extracted ion mobiligrams (EIMs) of derivatized GPG from WT and *CLN6* KO cells, shown alongside synthetic standards. GPG standards are d5-labeled. Each trace is independently normalized to its maximum peak intensity (100%) and therefore shows relative signal intensity rather than absolute abundance. **(C)** Proportion of endogenous GPG assigned to the *R,S* configuration in WT cells and the indicated single-, double-, and triple-knockout cell lines. **(D)** Experimental workflow for tracing the incorporation of d5-*R,S*-GPG into downstream lipid products in live cells. **(E)** Percentage of intracellular d5-GPG not assigned to the *R,S* configuration following 100 µM d5-*R,S*-GPG supplementation in the indicated cell lines, calculated as 100% minus the percentage of d5-*R,S*-GPG. The synthetic d5-*R,S*-GPG tracer was analyzed in parallel. The dashed line marks its approximately 6% stereoisomeric impurity baseline of the tracer. **(F)** Representative EICs of derivatized d5-LPG 18:1 from *CLN5* KO and *CLN5/CLN6* dKO cells following d5-*R,S*-GPG tracing. d5-LPG products containing unlabeled or deuterated monoacylglycerol (MAG) moieties were monitored using d0-MAG or d5-MAG fragment ions, respectively. Endogenous LPG 18:1 from *CLN5* KO cells is shown for comparison. **(G)** Relative proportions of tracer-derived d5-LPG 18:1 products distinguished by stereochemistry and MAG labeling in the cell lines shown in **(F)**. N.D., not detected. **(H)** Representative EICs of derivatized d5-BMP 18:1_18:1 from WT and *CLN6* KO cells following d5-*R,S*-GPG tracing with endogenous BMP 18:1_18:1 from *CLN6* KO and WT cells shown for comparison. **(I)** Representative EICs of derivatized d5-LPG 18:1 from CLN5 KO and CLN5/6 dKO cells following d5-*S,S*-GPG tracing. d5-LPG products were monitored using d0-MAG or d5-MAG fragment ions, respectively, with endogenous LPG 18:1 from *CLN5* KO cells shown for comparison. **(J)** Relative proportions of tracer-derived d5-*S,S*-LPG 18:1 products distinguished by MAG labeling in the cell lines shown in **(I)**. Data in **(C), (E)**, **(G),** and **(J)** are presented as mean ± SD of four biologically independent samples per group; points represent individual biological replicates. **(K)** Representative EICs of derivatized d5-BMP 18:1_18:1 from WT and CLN6 KO cells following d5-*S,S*-GPG tracing, with endogenous BMP 18:1_18:1 from WT cells shown for comparison. In **(F)**, **(H)**, **(I)**, and **(K)**, red and blue traces represent samples with and without tracer supplementation, respectively; gray traces show the endogenous lipids specified above. Dashed lines indicate the retention times of the indicated isomers. **(L)** Proposed pathways for the incorporation of d5-*R,S*-GPG into LPG and BMP through CLN6-dependent epimerization and subsequent reactions mediated by CLN8 and CLN5. Deuterium labels are shown in red. **(M)** Concentration-dependent changes in BMP 18:1_18:1 (top) and BMP 22:6_22:6 (bottom) following *S,S*-GPG supplementation in WT iNeurons and iNeurons derived from two independent *CLN6* KO clones. Abundances are expressed as fold changes relative to the corresponding untreated group (0 µM). Data are presented as mean ± SD. *n* = 3 independent biological samples. Panel D was created using BioRender (https://biorender.com).

To determine the stereochemistry of endogenous GPG, we developed a strategy combining chemical derivatization with high-resolution trapped ion mobility spectrometry (TIMS) to resolve these epimers (**fig. S3A**). By comparing the ion mobilities of endogenous GPG with synthetic deuterated *R,S*-GPG and *S,S*-GPG standards, we detected both epimers in WT cells, whereas the peak that represents *S,S*-GPG and *R,R*-GPG was absent in *CLN6* knockout cells (**Fig. 3B**). Although TIMS cannot distinguish the enantiomers *R,R*-GPG and *S,S*-GPG, it clearly separates these species from the *R,S*-GPG, enabling quantification of the *R,S*-GPG fraction, which is used in the following section.

To determine whether epimerization of *R,S*-GPG depends on other NCL genes involved in BMP biosynthesis, we quantified the proportion of endogenous *R,S*-GPG across cell lines deficient in different combinations of NCL genes (**Fig. 3C**). In addition to the previously established *CLN5* KO cells (*21*), we generated *CLN8* KO cells (**fig. S3B**), which showed depleted BMPs (**fig. S3C**). All cell lines lacking *CLN6*, including *CLN6* single-knockout, *CLN5/CLN6* and *CLN6/CLN8* double-knockout, and *CLN5/CLN6/CLN8* triple-knockout cells, contained predominantly *R,S*-GPG. In contrast, *R,S*-GPG accounted for approximately half of endogenous GPG in cells retaining the CLN6 protein (**Fig. 3C**).

To further assess the role of CLN6 in GPG epimerization and *S,S*-BMP generation, we performed isotope-tracing experiments by supplementing live cells with d5-*R,S*-GPG, which has deuterium labels on the *R*-glycerol (**Fig. 3D**). The synthetic tracer contained approximately 6% stereoisomeric impurities (**Fig. 3E**). Apparent isomerization exceeded this baseline in WT and *CLN8* knockout cells, reaching approximately 8% and 13%, respectively, but remained at or below baseline in *CLN6* knockout and *CLN6/CLN8* double-knockout cells (**Fig. 3E**), consistent with the requirement for CLN6, but not CLN8, in GPG epimerization. These measurements were done while cells are still incubated with excess d5-*R,S*-GPG hence the low percentage of detected isomerization product.

We next examined the stereochemical fate of the d5-*R,S*-GPG tracer during its incorporation into d5-LPG. As CLN5 quickly consumes LPG to synthesize BMP (*21*), we used *CLN5* KO cells to trap LPG from further conversion to determine its stereochemical configuration. When monitoring the d5-LPG 18:1 products, in *CLN5* KO cells, d5-*R,S*-GPG yielded predominantly d5-*S,S*-LPG 18:1, with the major chromatographic peak corresponding to the *sn*-2 position (**Fig. 3F**). The deuterium label was almost equally present on either the acylated or the nonacylated glycerol moiety, determined by a labeled (d5-MAG) or unlabeled (d0-MAG) monoacylglycerol moiety in MS/MS fragmentation, respectively. A minor fraction of tracer-derived d5-LPG 18:1 had the *S,R* configuration with acyl chain on d0-*S*-glycerol. In contrast, tracer-derived d5-LPG in *CLN5/CLN6* double-knockout cells consisted almost entirely of d5-*S,R*-LPG with an unlabeled MAG moiety, whereas both labeling forms of *S,S*-LPG were undetectable (**Fig. 3F**). Of note, the stereochemistry and regiochemistry of the d5-LPG 18:1 products are aligned with the endogenous LPG 18:1 from *CLN5* KO cells. Quantification of the relative proportions of these isomers in *CLN5*-knockout and *CLN5/CLN6* double-knockout cells confirms this shift in stereochemical composition (**Fig. 3G**).

Tracing into BMP revealed that d5-*R,S*-GPG gave rise to d5-*S,S*-BMP 18:1_18:1, predominantly the 2,2′ isomer, in WT cells, but not in *CLN6* knockout cells (**Fig. 3H**). Together with the LPG labeling patterns, these results indicate a pathway in which CLN6-dependent generation of *S,S*-GPG precedes the formation of *S,S*-LPG and BMP.

We then asked whether supplying *S,S*-GPG could bypass the requirement for CLN6. Following supplementation with d5-*S,S*-GPG, both *CLN5* knockout and *CLN5/CLN6* double-knockout cells produced predominantly d5-*S,S*-2-LPG 18:1 (**Fig. 3I**), with the label present approximately equally on either glycerol moiety (**Fig. 3J**). Moreover, d5-*S,S*-GPG was incorporated into d5-*S,S*-2,2′-BMP 18:1_18:1 in both WT and *CLN6* knockout cells (**Fig. 3K**). Thus, cells lacking CLN6 retained the capacity to convert *S,S*-GPG into downstream *S,S*-LPG and BMP.

These results can be explained by our model of d5-BMP biosynthesis from the d5-*R,S*-GPG tracer (**Fig. 3L**). In the absence of CLN6, d5-*R,S*-GPG retains the *R,S* stereochemistry and undergoes CLN8-mediated acylation at the *sn*-2 position of its unlabeled, *S*-glycerol moiety. This generates d5-*S,R*-2-LPG with an unlabeled monoacylglycerol moiety, which could be monitored by the d0-MAG fragment ion in MS/MS. Subsequent acylation by CLN5 produces the unstable d5-*R,S*-2,2′-BMP. In the presence of CLN6, d5-*R,S*-GPG first undergoes epimerization to d5-*S,S*-GPG. CLN8 can then acylate either *S*-glycerol moiety, generating d5-*S,S*-2-LPG with d5-MAG or d0-MAG moieties in approximately equal proportions. Subsequent CLN5-mediated acylation of d5-*S,S*-2-LPG yields d5-*S,S*-2,2′-BMP.

Consistent with this model, supplementation with exogenous *S,S*-GPG increased BMP abundance in a concentration-dependent manner in iNeurons derived from two independent *CLN6*-knockout clones, whereas WT iNeurons showed comparatively small increase above their baseline, consistent with abundant precursor in WT cells (**Fig. 3M & fig. S3D)**. These findings support the generation of *S,S*-GPG as a key CLN6-dependent step in BMP biosynthesis and show that supplying this intermediate can bypass the requirement for CLN6 in downstream lipid synthesis.

Finally, we asked whether supplying *S,S*-GPG could alleviate lipid accumulation associated with CLN6 deficiency using ARPE-19/HPV-16 cell model (ARPE-19 hereafter), a commonly used cell line to study the biochemical pathology in Batten disease (*32–34*). In two independent *CLN6* KO ARPE-19 clones (**fig. S3E**), *S,S*-GPG supplementation reduced filipin staining of cholesterol and GM2 immunofluorescence (**fig. S3F-I**). Thus, supplying this biosynthetic intermediate can bypass CLN6 function and mitigate the lipid-storage phenotype in CLN6-deficient cells.

### CLN6 is a calcium-dependent GPG epimerase

Based on these results, we wondered if CLN6 is a GPG epimerase. We next purified CLN6 protein (**fig. S4A-B & E**). WT CLN6 protein incubated with d5-*R,S*-GPG showed no epimerization, which prompted us to test if a metal cofactor is needed (**Fig. 4A**). Indeed, supplementation with either Dulbecco’s Modified Eagle Medium (DMEM), which provides a range of metal ions, or calcium allowed the detection of GPG epimerization (**Fig. 4A**). Consistently, neither WT HEK293T cell lysate treated with BAPTA, a Ca^2+^ chelator, nor *CLN6* KO lysate showed GPG epimerization (**Fig. 4B)**. Purified CLN6 then showed a Ca^2+^-dependent epimerase activity, with an endpoint EC50 of approximately 900 µM Ca^2+^ (**Fig. 4C**). Furthermore, the amount of epimerized product increased in both a time– and substrate concentration-dependent manner (**Fig. 4D-E**). Of importance, the purified patient-derived H127Y variant (**fig. S4C-E**) exhibited no detectable epimerase activity (**Fig. 4F**).

**Figure 4.**
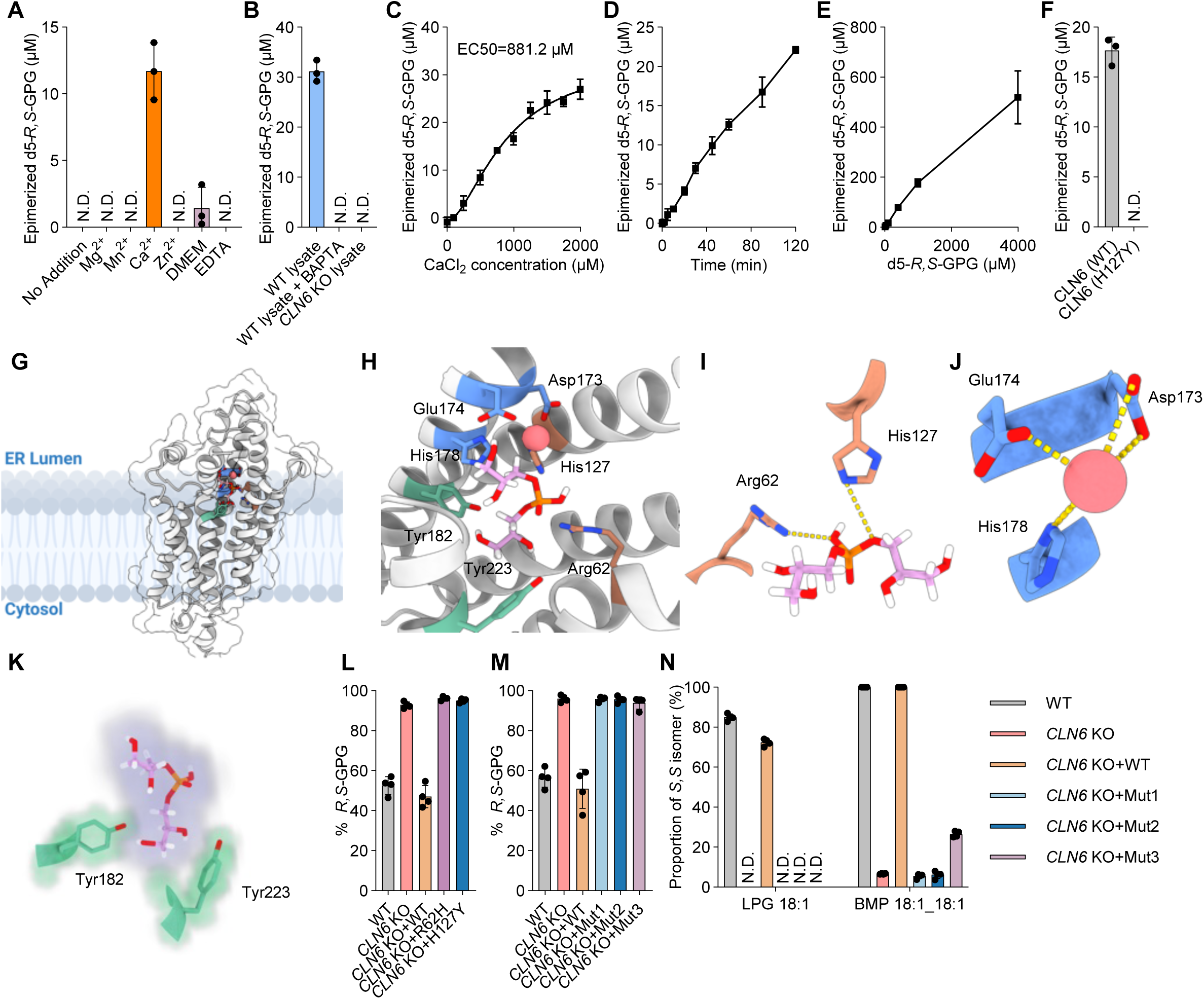
CLN6 is a calcium-dependent GPG epimerase. **(A)** Epimerization of 100 µM d5-*R,S*-GPG by purified CLN6 in the presence of the indicated metal ions, DMEM, or EDTA, compared with a control without metal ion supplementation. *n* = 3 biologically independent samples per condition. **(B)** Epimerization of 100 µM d5-*R,S*-GPG following overnight incubation with WT cell lysates, WT lysates treated with the Ca^2+^ chelator BAPTA, and *CLN6* KO cell lysates. *n* = 3 independent biological samples per condition. **(C)** Epimerization of 100 µM d5-*R,S*-GPG by purified CLN6 at increasing CaCl_2_ concentrations after 120 min. *n* = 2 at 1000 µM CaCl_2_ and *n* = 3 other concentrations. Data were fitted using a four-parameter logistic model, yielding an EC50 of 881.2 µM for CaCl□. **(D)** Time course of epimerization of 100 µM d5-*R,S*-GPG by purified CLN6 in the presence of 500 µM CaCl_2_. *n* = 3 for 0 min and *n* = 4 for other incubation times. **(E)** Epimerization of d5-*R,S*-GPG by purified CLN6 at increasing substrate concentrations in the presence of 500 µM CaCl_2_ for 120 min. *n* = 3 independent biological samples. **(F)** Epimerization of 100 µM d5-*R,S*-GPG by purified WT CLN6 or the patient-derived H127Y variant in the presence of 500 µM CaCl_2_ for 120 min. *n* = 3 independent biological samples. In **(A–F)**, data are presented as mean ± SD, and epimerization is expressed as the concentration of isomerized d5-GPG. N.D., not detected. **(G)** Structural model of CLN6 with docked *R,S*-GPG and Ca^2+^, generated using Schrödinger software. The predicted orientation relative to the ER lumen and cytosol is indicated. **(H)** Enlarged view of the predicted GPG– and Ca^2+^-binding pocket. **(I)** Predicted interactions of GPG with R62 and H127, residues affected by the patient-derived R62H and H127Y variants. **(J)** Predicted Ca^2+^ coordination by D173, E174, and H178. Ca^2+^ is shown as a pink sphere. **(K)** Positions of Y182 and Y223 relative to GPG in the predicted substrate-binding pocket. **(L)** Proportion of endogenous GPG assigned to the *R,S* configuration in WT cells, *CLN6* KO cells, and KO cells re-expressing WT CLN6 or patient variants. **(M and N)** Proportion of endogenous GPG assigned to the *R,S* configuration **(M)** and the *S,S* isomers of endogenous LPG 18:1 and BMP 18:1_18:1 **(N)** in WT cells, *CLN6* KO cells, and KO cells re-expressing WT CLN6 or the indicated mutants (Mut1–Mut3). In **(L–N)**, data are presented as mean ± SD. *n* = 4 biologically independent samples per group.

To identify residues potentially involved in Ca^2+^ binding and substrate recognition, we examined an AlphaFold-based structural model of CLN6 (**Fig. 4G & H**). The model predicted interactions between *R,S*-GPG and R62 and H127, the residues affected by the patient-derived R62H and H127Y variants (**Fig. 4I**) and suggested a putative Ca^2+^-binding site involving D173, E174, and H178 (**Fig. 4J**). Y182 and Y223 were also positioned near GPG in the predicted substrate-binding pocket (**Fig. 4K**). Interestingly, these residues are facing the ER lumen, consistent with the higher concentration of Ca² in the ER. Re-expression of WT CLN6 in *CLN6*-knockout cells restored the proportion of *R,S*-GPG to approximately WT levels, whereas neither R62H nor H127Y restored GPG stereochemistry (**Fig. 4L**).

We next tested the functional importance of the other residues by expressing CLN6 variants in *CLN6* KO cells (**fig. S4F-H**). Neither the D173A/E174A/H178A (Mut1) nor the D173Q/E174Q/H178A (Mut2) restored GPG stereochemistry (**Fig. 4M**), BMP abundance (**fig. S4I**), or the predominantly *S,S* configuration of LPG and BMP (**Fig. 4N**). The Y182A/Y223A (Mut3) double mutant similarly failed to restore GPG stereochemistry (**Fig. 4M**), BMP abundance (**fig. S4I**) and LPG stereochemistry and only partially increased *S,S*-BMP (**Fig. 4N**). Altogether, these results confirm that CLN6 is necessary and sufficient for the epimerization of GPG and characterize the essential residues required for its enzyme activity.

### CLN6-deficient mice and NCL6 patients exhibit altered stereochemistry of GPG, LPG and BMP

We examined the stereochemistry of GPG, LPG, and BMP in the *Cln6*^nclf^ mouse model (*35*). Consistent with our in vitro data, *R,S*-GPG was the predominant isomer in *Cln6*^nclf/nclf^-derived brain and lung tissues compared to approximately 70% in the heterozygous controls *Cln6*^wt/nclf^ (**Fig. 5A**). BMP levels showed significant decrease in homozygous mutants compared to carriers in both brain (**fig. S5A**) and lung tissues (**fig. S5B**). Heterozygous controls have almost exclusively *S,S*-BMP, whereas *S,S*-BMP accounted for only approximately 20% of brain BMP and 10% of lung BMP in homozygous mutants (**Fig. 5B**). Although LPG levels are similar between heterozygous and homozygous mice in both brain (**fig. S5C**) and lung tissues (**fig. S5D**), *S,S*-LPG is the predominant LPG in control brain and accounted for approximately 50% of lung LPG but was undetectable in either tissue from homozygous mutants (**Fig. 5C**). In *CLN6*-knockout human iNeurons, *R,S*-GPG was also the predominant GPG stereoisomer compared to approximately 50% of GPG in WT cells (**Fig. 5D**). Likewise, *S,S*-LPG and *S,S*-BMP predominated in WT iNeurons but were undetectable in *CLN6*-knockout iNeurons (**Fig. 5E and F**).

**Figure 5.**
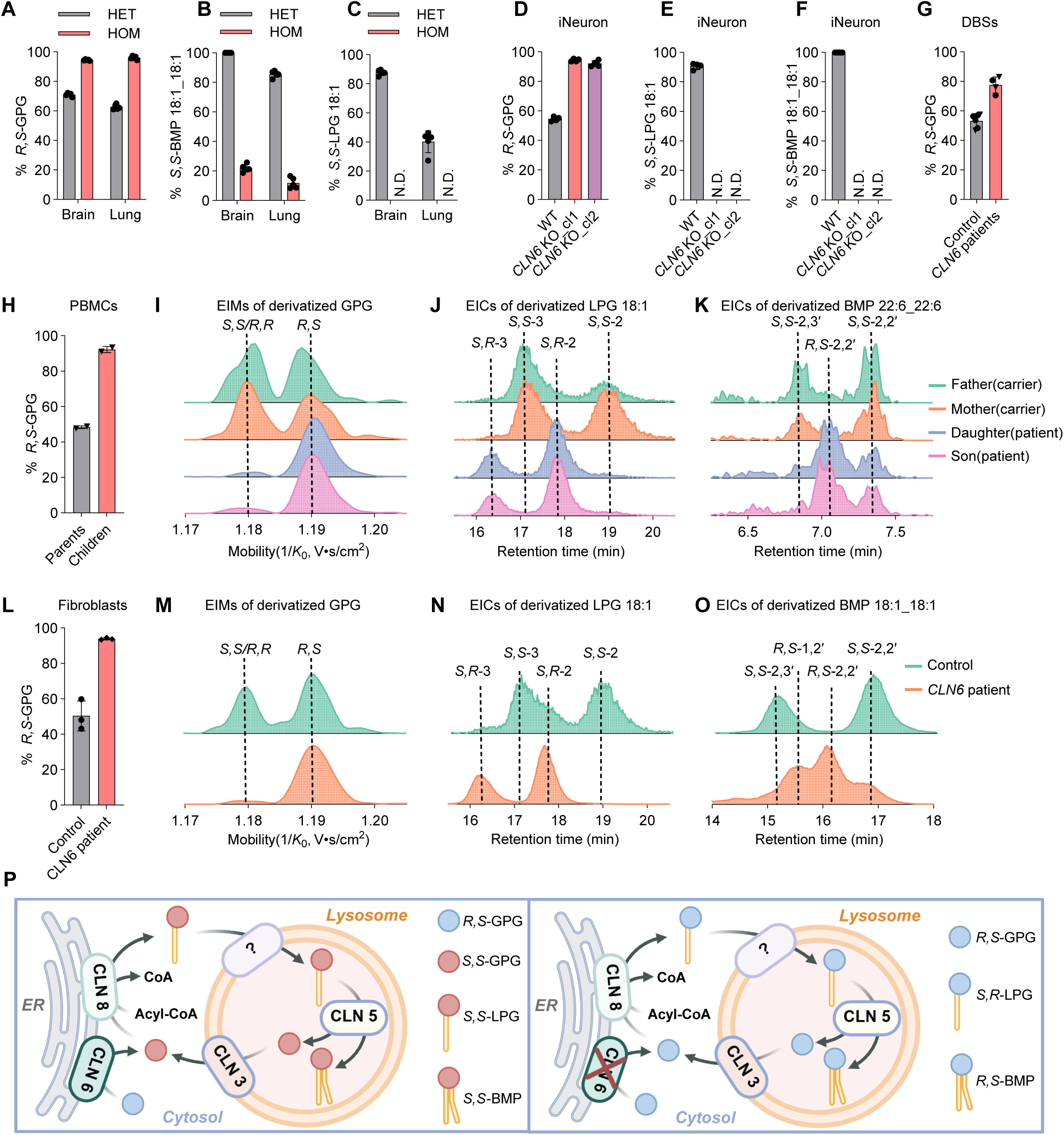
CLN6-deficient mice and NCL6 patients exhibit altered stereochemistry of GPG, LPG and BMP. (**A-C**) Proportions of *R,S*-GPG **(A)**, *S,S*-BMP 18:1_18:1 **(B)**, and *S,S*-LPG 18:1 **(C)** in brain and lung tissues from heterozygous *Cln6* carrier mice (HET; WT/*nclf*) and homozygous mutant mice (HOM; *nclf/nclf*). N.D., not detected. Each genotype measurement represents *n* = 5 biologically independent samples. **(D-F)** Proportion of *R,S*-GPG **(D)**, *S,S*-LPG 18:1 **(E)**, and *S,S*-BMP 18:1_18:1 **(F)** in WT iNeurons and iNeurons derived from two independent *CLN6* KO clones. Each genotype measurement represents *n* = 4 biologically independent samples. **(G)** Proportion of *R,S*-GPG in DBS from patients with CLN6 disease and healthy controls and heterozygous carrier parents (dots denote individuals from Family One and their controls and triangles denote individuals from Family Two). **(H)** Proportion of *R,S*-GPG in PBMCs from two heterozygous carrier parents and their two affected children from Family Two. **(I–K)** EIMs of derivatized GPG **(I)** and extracted ion chromatograms (EICs) of derivatized LPG 18:1 **(J)** and BMP 22:6_22:6 **(K)** from PBMCs of the family shown in **(H)**. Green, orange, blue, and magenta traces correspond to the father, mother, daughter, and son, respectively. **(L)** Proportion of *R,S*-GPG in fibroblasts from healthy controls and a patient with CLN6 disease of Family Three. Dots represent three healthy controls; diamonds represent three technical replicates of the patient-derived sample. **(M–O)** Representative EIMs of derivatized GPG **(M)** and EICs of derivatized LPG 18:1 **(N)** and BMP 18:1_18:1 **(O)** from control and patient-derived fibroblasts. Green and orange traces represent healthy control and the patient samples, respectively. In **(I–K)** and **(M–O)**, each trace is independently normalized to its maximum peak intensity (100%) and therefore shows relative signal intensity rather than absolute abundance. In **(J)**, **(K)**, **(N)**, and **(O)**, dashed lines indicate the retention times of the indicated LPG and BMP isomers. **(P)** Proposed model of CLN6-dependent control of BMP stereochemistry. With functional CLN6 (left), GPG epimerization supplies *S,S*-GPG for downstream LPG and BMP synthesis mediated by CLN8 and CLN5. In the absence of functional CLN6 (right), *R,S*-GPG predominates, favoring the formation of *S,R*-LPG and *R,S*-BMP. CLN3, required for GPG efflux, and an unidentified LPG transport step are included in the proposed model. Panel P was created using BioRender (https://biorender.com).

Dried blood spots (DBSs) from NCL6 patients collected at German (Family One) and US sites (Family Two) showed that *R,S*-GPG represents 80% of the total GPG compared to approximately 50% in healthy controls for Family One and heterozygous carrier parents for Family Two (**Fig. 5G**). BMP levels of NCL6 patients also showed a general trend of decrease compared to healthy controls or carrier parents (**fig. S5E&F**), while LPG did not show significant changes (**fig. S5G&H**).

We further examined peripheral blood mononuclear cells (PBMCs) from a family with two heterozygous carrier parents and two children with NCL6 (Family Two). *R,S*-GPG predominated in both affected children but accounted for only ∼50% of GPG in their carrier parents (**Fig. 5H**), which was also evident by distinctly different mobiligrams (**Fig. 5I**). LPG abundance was variable between affected children and their carrier parents (**fig. S5I**), however, subsequent stereochemistry measurement of LPG (**Fig. 5J**) confirmed the loss in *S,S* stereochemistry. While severe depletion of BMP 18:1_18:1 in patient samples hindered our ability to measure its stereochemistry (**fig. S5J**), we observed a distinct shift in the chiral chromatogram of BMP 22:6_22:6 (**Fig. 5K**), indicating a loss of its canonical stereochemistry. To overcome this limitation, we examined fibroblasts from an NCL6 patient (Family Three) and healthy controls. Like PBMCs, the patient showed predominantly *R,S*-GPG compared to ∼50% in healthy controls (**Fig. 5L & M**). Consistently, LPG (**Fig. 5N**) and BMP (**Fig. 5O**) lacked *S,S* stereochemistry. Altogether, our data confirm that CLN6 is required for establishing *S,S* stereochemistry of GPG, LPG, and BMP in mice and humans.

## Discussion

BMP is key to lysosomal lipid catabolism and trafficking (*13*) and its dyshomeostasis is associated with neurodegeneration and aging (*6, 36, 37*). BMP function depends on its stability in the lysosome and requires both its unique *S,S* stereochemistry and 2,2′ regiochemistry to resist phospholipase activity (*30*). More than half a century ago, this unique stereochemistry was assigned (*18, 19*), however, its molecular machinery remained elusive. Our findings identify GPG epimerization as a critical step in establishing the distinctive *S,S* stereochemistry of BMP defining a pathway in which stereochemical inversion happened ahead of acylation: *R,S*-GPG → *S,S*-GPG → *S,S*-2-LPG → *S,S*-2,2′-BMP. In this pathway, CLN6 catalyzes GPG epimerization from *R,S* to *S,S*, CLN8 acylates *S,S*-GPG to generate *S,S*-LPG (*24*), and CLN5 converts *S,S*-LPG to *S,S*-BMP (*21, 38*). Several observations support this pathway as a major route to BMP. First, *CLN8*-knockout cells contain almost no detectable LPG or BMP (*23, 24*). Second, the stereochemical identities of GPG and LPG in cells lacking CLN6 or CLN8 are consistent with their proposed positions in the pathway. Third, the requirement for CLN6 in *S,S*-GPG production complements the stereoselectivity of CLN8 for acylation of an *S*-configured glycerol moiety. Together, these findings place three Batten disease genes CLN6, CLN8, and CLN5 at successive steps in *S,S*-BMP biosynthesis and provide a shared biochemical explanation for the BMP deficiency associated with loss of these NCL proteins.

Our data demonstrate that CLN6 is a Ca^2+^-dependent epimerase. An independent study using complementary approaches also identifies CLN6 as a GPG epimerase [Sheokand et al., *bioRxiv*, 2026]. Our findings support a model in which CLN3-dependent GPG efflux from lysosomes (*28*) is coupled to ER-associated precursor processing and lysosomal BMP synthesis (**Fig. 5P**). In this model, CLN6 converts *R,S*-GPG to *S,S*-GPG, which CLN8 acylates to form *S,S*-LPG. Subsequent delivery of *S,S*-LPG to lysosomes through an unidentified transport mechanism enables CLN5-dependent synthesis of *S,S*-BMP. Without functional CLN6, *R,S*-GPG remains available for CLN8-mediated acylation, yielding *S,R*-LPG and, following CLN5-dependent conversion, *R,S*-BMP, although at much lower efficiency as indicated by the net depletion of BMP. ER is the major intracellular Ca^2+^ store, maintaining a luminal free homeostatic Ca^2+^ concentration of approximately 500 µM (*39*). Notably, CLN6 shows little activity at Ca^2+^ concentrations below 100 µM, suggesting that its catalytic activity is specifically adapted to the Ca^2+^-rich environment of the ER lumen. Therefore, persistent depletion of ER Ca^2+^ could have two coordinated effects on lysosomal homeostasis: suppression of CLN6 catalytic activity and impairment of lysosomal Ca^2+^ replenishment. Reduced lysosomal Ca^2+^ could weaken local signals that support autophagosome-lysosome fusion (*40*), lysosomal acidification (*41*), and calcineurin-TFEB mediated lysosomal biogenesis (*42*).

Finally, our findings support precursor supplementation as a potential strategy for restoring BMP synthesis. Increasing BMP has been reported to lower cellular cholesterol accumulation in lysosomal storage diseases, and we have recently shown that PLA2G15 knockout increases cellular BMP levels and ameliorates NPC1 pathology (*30*). The predominance of *R,S*-GPG in NCL6 patient-derived cells and DBS suggests a potential biochemical marker of CLN6 dysfunction. Measuring the *R,S* proportion of GPG could complement genetic testing, and measuring *S,S*-LPG and *S,S*-BMP could provide an interventional readout for the responsiveness to treatments activating CLN6 variants. Future studies should also elucidate the origin of *R,S*-GPG, potentially in the mitochondria, and the mechanisms governing *S,S*-LPG trafficking between its sites of synthesis in the ER and downstream conversion in the lysosome, thereby clarifying how BMP synthesis is coordinated across cellular compartments.

## Supplementary Figures

**Figure S1.**
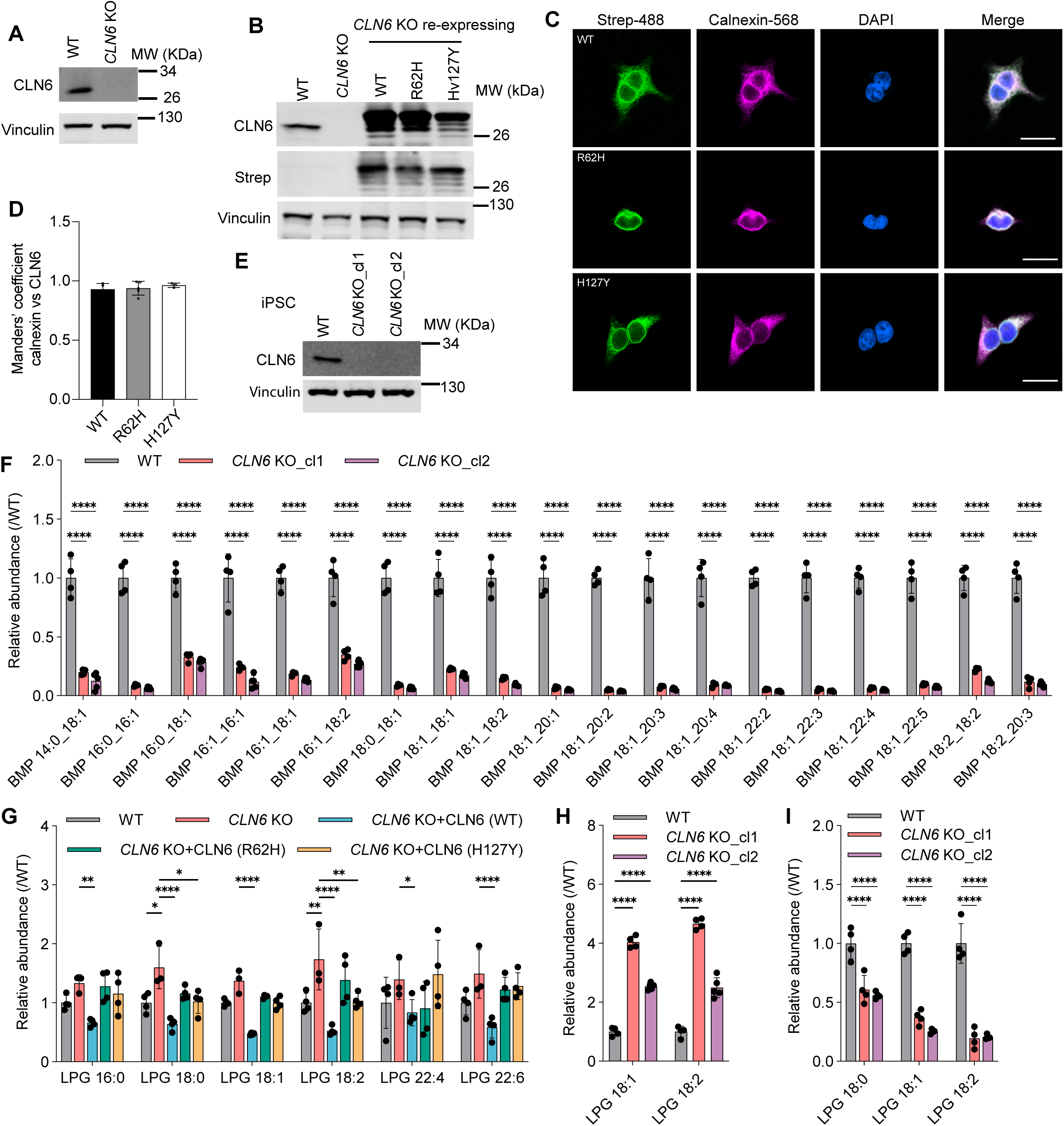
Validation of CLN6-deficient cell models. **(A)** Immunoblot of CLN6 and vinculin in WT and *CLN6*-KO HEK293T cells. **(B)** Immunoblot of CLN6, Twin-Strep-tagged CLN6 (detected with anti-Strep antibody) and vinculin in WT, *CLN6* KO and *CLN6*-KO HEK293T cells expressing WT CLN6 or the patient R62H and H127Y variants, C-terminus of each is fused to Twin-Strep tag. **(C)** Representative immunofluorescence images of *CLN6*-KO HEK293T cells expressing Strep-tagged WT CLN6, R62H, or H127Y. Strep-tagged CLN6 is shown in green, the ER marker calnexin in magenta, and nuclei stained with DAPI in blue. Merged images are shown on the right. Scale bars, 20 µm. **(D)** Quantification of Manders’ colocalization coefficient between CLN6-2XStrep and calnexin from **(C)**. Each dot represents one imaging field. Data are presented as mean ± SD, *n* = 4 fields for WT, 5 fields for R62H, and 3 fields for H127Y. **(E)** Immunoblot of CLN6 and vinculin in WT and *CLN6* knockout iPSCs. **(F)** Targeted analysis of BMP species in WT iPSCs and two independent *CLN6*-KO clones (cl1 and cl2). Data are presented as mean ± SD; *n* = 5 biologically independent samples for the KO clone 2 and *n* = 4 for each remaining group. **(G)** Targeted lipidomics analysis of LPG species in WT HEK293T cells, *CLN6*-KO cells, and KO cells re-expressing WT CLN6 or patient-derived variants, R62H or H127Y. Data are presented as mean ± SD; *n* = 3 biologically independent samples for the KO group and *n* = 4 for each remaining group. **(H and I)** Targeted analysis of LPG species in WT cells and two independent *CLN6*-KO clones (cl1 and cl2), measured in iPSCs **(H)** and iNeurons **(I)**. Data are presented as mean ± SD; *n* = 5 biologically independent samples for iPSC KO clone 2 and *n* = 4 for each remaining group. In **(F–I)**, the abundance of each lipid species was first normalized to endogenous POPC and then normalized to its mean abundance in the corresponding WT group. Statistical significance in **(F–I)** was assessed using two-way ANOVA; \**P* < 0.05, \*\**P* < 0.01, and \*\*\*\**P* < 0.0001.

**Figure S2.**
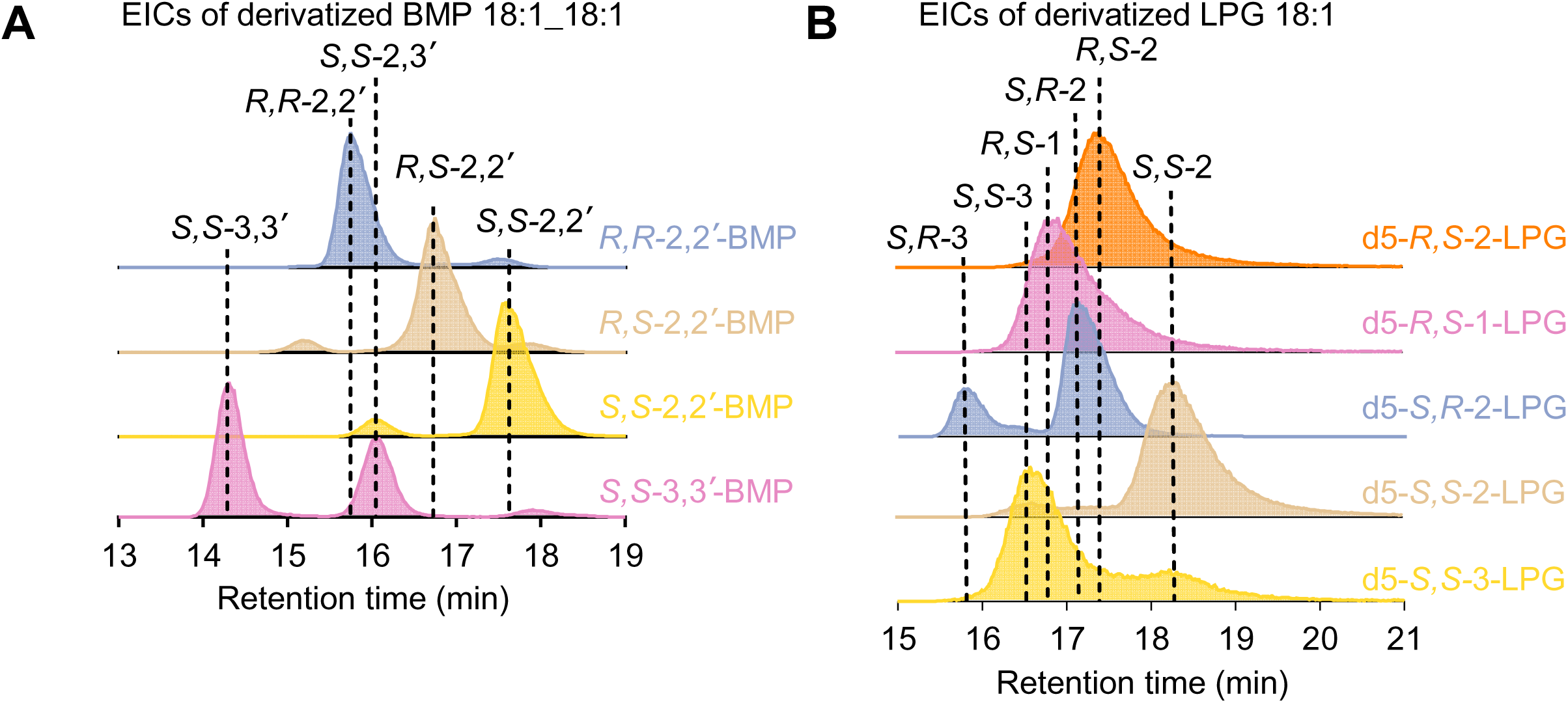
Development of chiral chromatographic methods to resolve the stereochemistry and regiochemistry of BMP and LPG. **(A)** Representative extracted ion chromatograms (EICs) of derivatized synthetic BMP 18:1_18:1 standards: *R,R*-2,2′-BMP, *R,S*-2,2′-BMP, *S,S*-2,2′-BMP, and *S,S*-3,3′-BMP, shown in blue, beige, yellow, and magenta, respectively. Impurities are found in some standards because of common acyl migration. **(B)** Representative EICs of derivatized synthetic deuterium-labeled LPG 18:1 standards: d5-*R,S*-2-LPG, d5-*R,S*-1-LPG, d5-*S,R*-2-LPG, d5-*S,S*-2-LPG, and d5-*S,S*-3-LPG, shown in orange, magenta, blue, beige, and yellow, respectively. Impurities are found in some standards because of common acyl migration. Each trace is independently normalized to its maximum peak intensity (100%). Dashed lines indicate the retention times of the assigned isomers. Traces are vertically offset for clarity.

**Figure S3.**
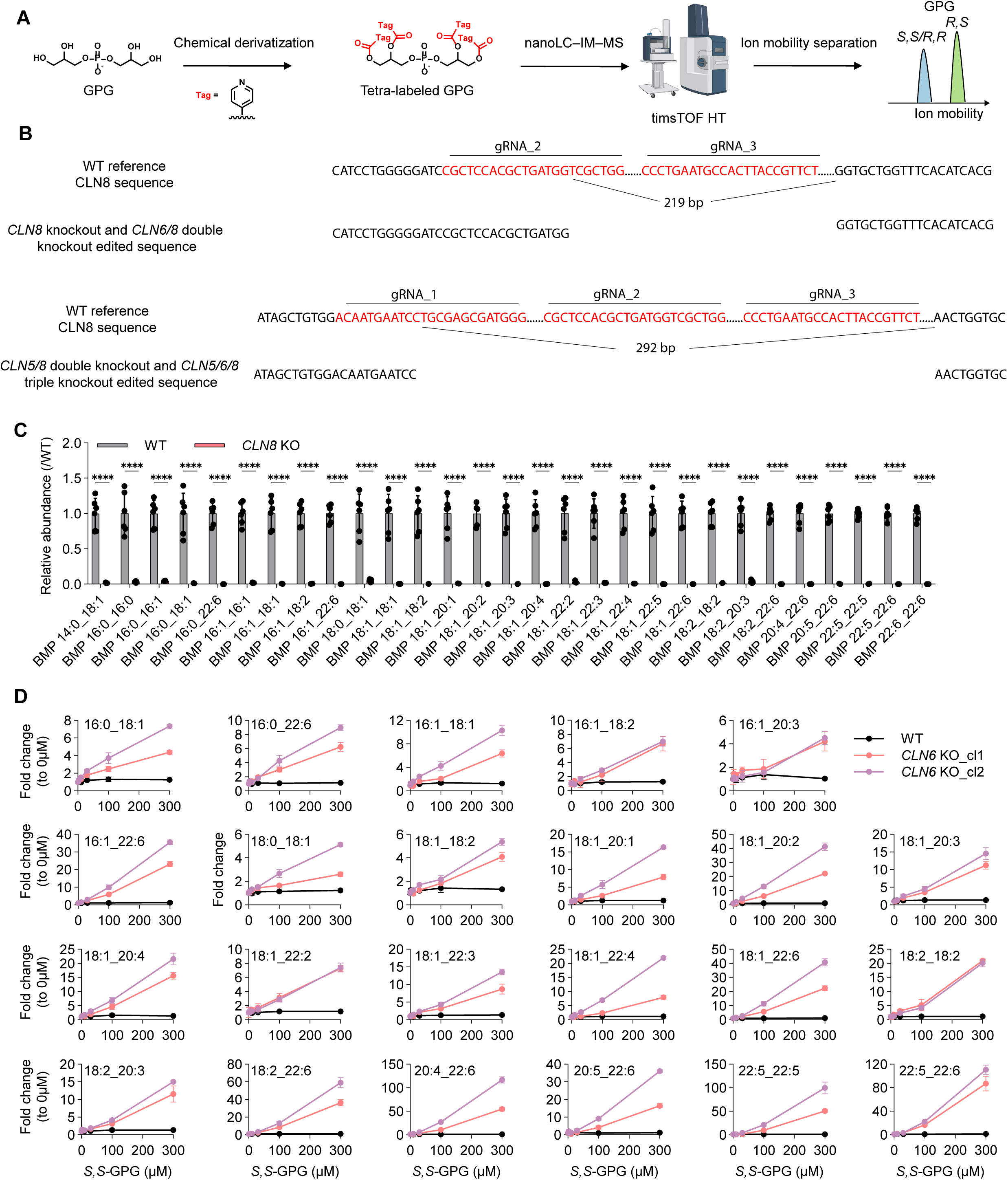
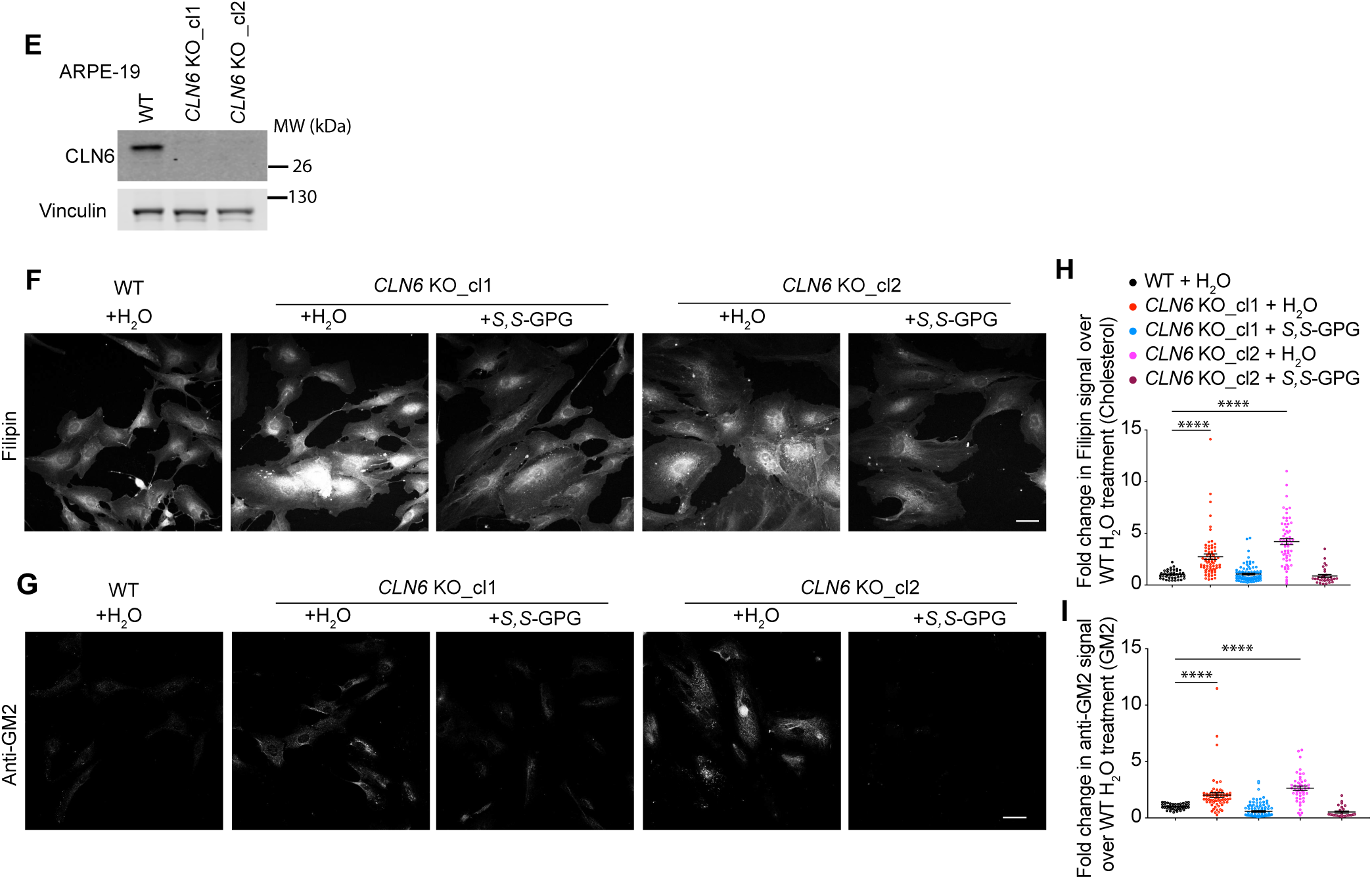
Method for GPG stereochemistry, validation of CLN8-deficient cells, and BMP responses to *S,S*-GPG supplementation. **(A)** Workflow for analyzing GPG stereochemistry by chemical derivatization and nanoLC–ion mobility–mass spectrometry using a timsTOF HT instrument. **(B)** CRISPR-Cas9 mediated deletion of CLN8 in single and combinatorial knockout cell lines. Alignment of the wild-type (WT) *CLN8* reference sequence with edited sequences in *CLN8* knockout and *CLN6/8* double-knockout cells (top), and *CLN5/8* double-knockout and *CLN5/6/8* triple-knockout cells (bottom). The edited alleles contain 219-bp and 292-bp disrupted nucleotides, respectively. gRNA target sequences are highlighted in red, with corresponding gRNAs indicated above. Flanking sequences retained in the edited alleles are shown in black. Ellipses indicated omitted sequence. *CLN5*/8 double-knockout was generated on a previously reported *CLN5* knockout (*21*), and *CLN5*/6/8 triple-knockout was generated based on the *CLN5*/8 double-knockout. *CLN6*/8 double-knockout was generated based on the *CLN8* knockout. **(C)** Targeted analysis of BMP species in WT and *CLN8*-knockout (KO) cells. The abundance of each BMP species was first normalized to endogenous POPC and then normalized to its mean abundance in the WT group. Data are presented as mean ± SD of *n* = 6 biologically independent samples. Statistical significance was assessed using two-way ANOVA. \*\*\*\**P* < 0.0001. **(D)** Concentration-dependent changes in indicated BMP abundances following supplementation with exogenous *S,S*-GPG in WT iNeurons and iNeurons derived from two independent *CLN6*-KO clones (cl1 and cl2). Abundances are expressed as fold changes relative to the corresponding untreated group (0 µM). Black, pink, and purple curves represent WT, KO clone 1, and KO clone 2, respectively. Data are presented as mean ± SD. *n* = 3 independent biological samples. **(E)** Immunoblot validation of CLN6 loss in two independent *CLN6*-knockout ARPE-19 clones (cl1 and cl2), with WT cells included as a control. Vinculin serves as a loading control. **(F and G)** Representative fluorescence images of vehicle-treated WT ARPE-19 cells and two independent *CLN6*-knockout clones (*CLN6* KO_cl1 and *CLN6* KO_cl2) treated with vehicle (H_2_O) or 400 µM *S,S*-GPG for 10 days. Cells were stained with filipin to visualize cholesterol **(F)** or a mouse anti-GM2 antibody to detect ganglioside GM2 **(G)**. Scale bars, 20 µm. **(H and I)** Quantification of filipin and GM2 fluorescence from **(F)** and **(G)**. Fluorescence intensity was normalized to cell area and expressed relative to the mean of vehicle-treated WT cells. Each dot represents one cell, and n denotes the total number of cells analyzed from 4-5 randomly selected imaging fields per condition. For filipin, *n* = 39 for H_2_O treated WT cells, 68 for H_2_O treated *CLN6* KO_cl1, 89 for *S,S*-GPG *CLN6* KO_cl1, 60 for H_2_O treated *CLN6* KO_cl2, and 31 for *S,S*-GPG treated *CLN6* KO_cl2. For GM2, *n* = 41 for H_2_O treated WT cells, 64 for H_2_O treated *CLN6* KO_cl1, 100 for *S,S*-GPG *CLN6* KO_cl1, 45 for H_2_O treated *CLN6* KO_cl2, and 27 for *S,S*-GPG treated *CLN6* KO_cl2. Statistical significance was assessed using one-way ANOVA; \*\**P* < 0.01 and \*\*\*\**P* < 0.0001. Panel A was created using BioRender (https://biorender.com).

**Figure S4.**
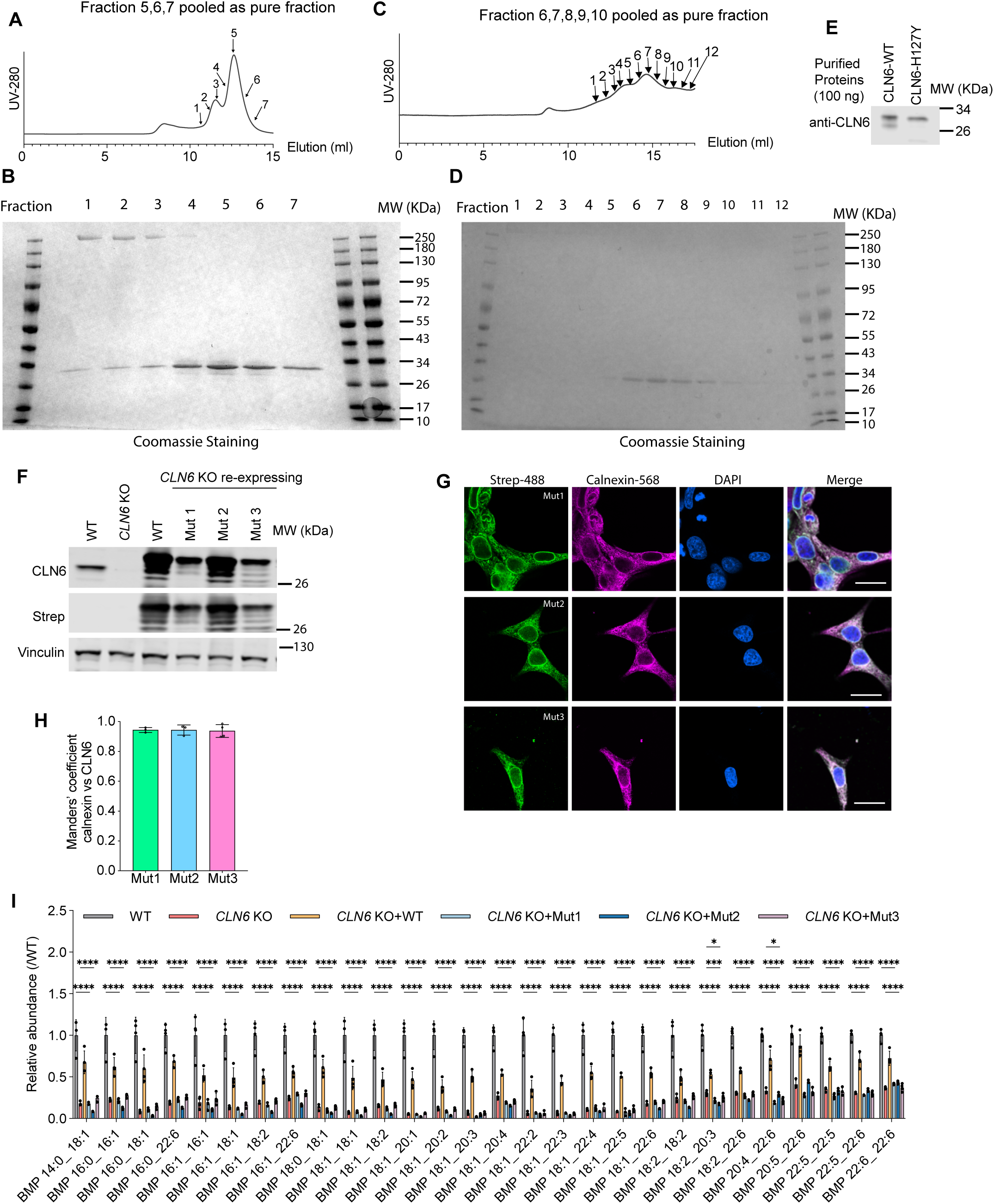
Purification of CLN6 and functional characterization of variants targeting putative Ca^2+^– and GPG-binding residues. (**A and B**) Size-exclusion chromatography profile of WT CLN6-2×Strep purified from Expi293 cells **(A)** and Coomassie-stained SDS–PAGE analysis of the indicated fractions **(B)**. Numbers above the chromatogram indicate collected fractions. Fractions 5-7 were pooled. **(C and D)** Size-exclusion chromatography profile of the H127Y variant CLN6-2×Strep purified from Expi293 cells **(C)** and Coomassie-stained SDS–PAGE analysis of the indicated fractions **(D)**. Fractions 6-10 were pooled. **(E)** Immunoblot analysis of purified WT CLN6 and H127Y using an anti-CLN6 antibody. 100 ng of protein was loaded. **(F)** Immunoblot validation of WT CLN6 and the indicated variants re-expressed in *CLN6*-KO cells, using anti-CLN6 and anti-Strep antibodies. WT and KO cells are included as controls. Vinculin serves as a loading control. Mut1, D173A/E174A/H178A; Mut2, D173Q/E174Q/H178A; Mut3, Y182A/Y223A. **(G)** Representative immunofluorescence images of *CLN6*-knockout (KO) cells expressing the indicated Strep-tagged CLN6 variants. Strep-tagged CLN6 is shown in green, the ER marker calnexin in magenta, and nuclei stained with DAPI in blue. Merged images are shown on the right. Scale bars, 20 µm. **(H)** Quantification using Manders’ colocalization coefficient of CLN6-2×Strep and calnexin overlap in Mut 1, Mut2, and Mut3 in **(G)**. Each dot represents one imaging field; *n* = 3 fields for Mut1 and Mut2, *n* = 4 fields for Mut3. **(I)** Targeted analysis of BMP species in WT cells, *CLN6*-KO cells, and KO cells re-expressing WT CLN6 or the indicated variants. The abundance of each BMP species was first normalized to endogenous POPC and then normalized to its mean abundance in the WT group. Data are presented as mean ± SD. *n* = 4 independent biological samples. Statistical significance was assessed using two-way ANOVA. Asterisks indicate statistical significance: \**P* < 0.05, \*\*\**P* < 0.001, and \*\*\*\**P* < 0.0001.

**Figure S5.**
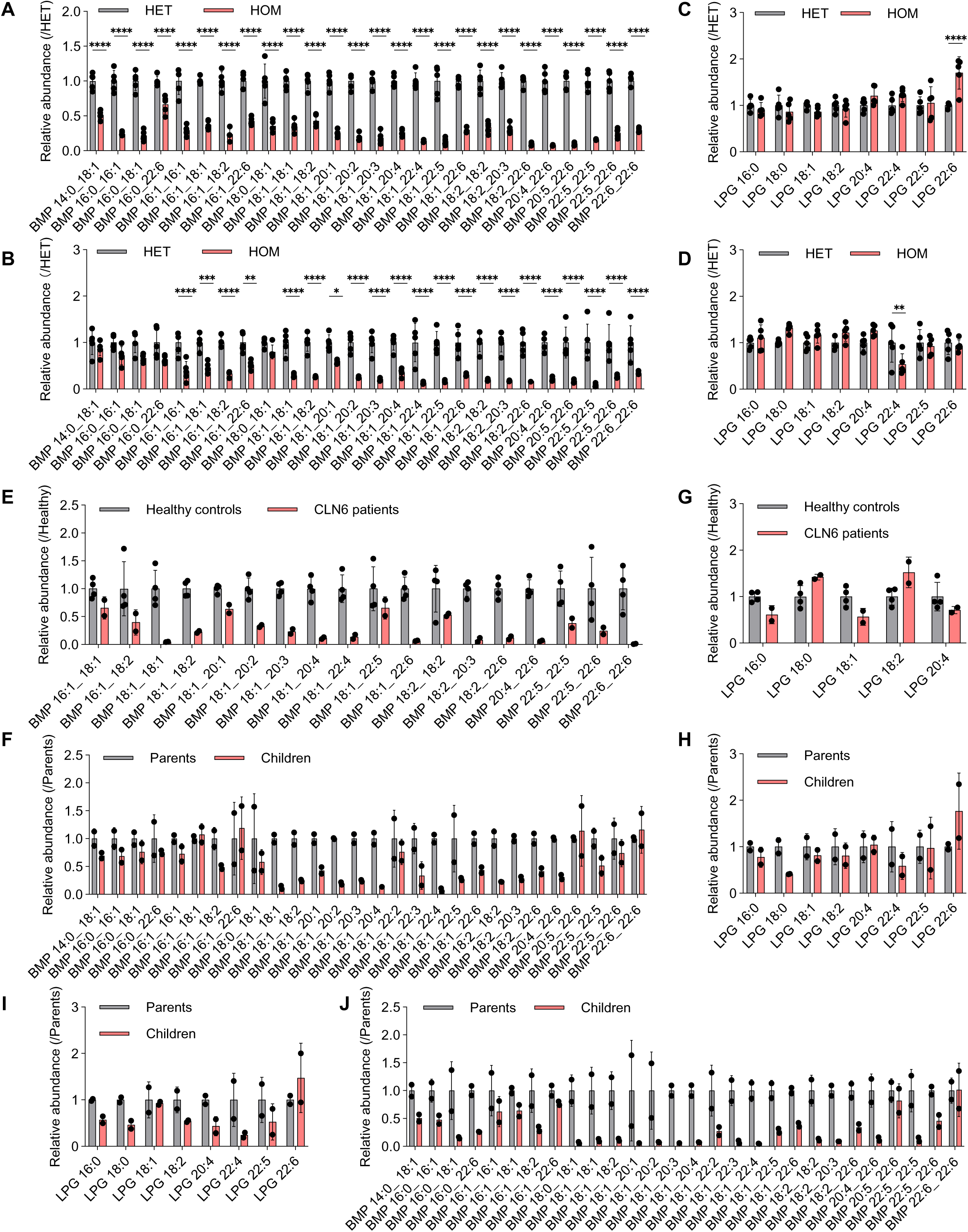
Quantitative analysis of BMP and LPG in mouse tissues and clinical samples associated with CLN6 deficiency. (**A–D**) Targeted analysis of BMP **(A and B)** and LPG **(C and D)** species in brain **(A and C)** and lung **(B and D)** tissues from heterozygous carrier (HET; *Cln6* WT/*nclf*) and homozygous mutant mice (HOM; *Cln6 nclf/nclf*). The abundance of each lipid species was first normalized to endogenous POPC and then normalized to its mean abundance in the corresponding HET tissue group. Gray and pink bars represent HET and HOM mice, respectively. Data are presented as mean ± SD. *n* = 5 independent biological samples. Asterisks indicate statistical significance: \**P* < 0.05, \*\**P* < 0.01, \*\*\**P* < 0.001, and \*\*\*\**P* < 0.0001. **(E and G)** Relative abundances of BMP **(E)** and LPG **(G)** species in dried blood spots (DBS) from two patients with CLN6 disease in Family One and four healthy controls. **(F and H)** Relative abundances of BMP **(F)** and LPG **(H)** species in DBS from two heterozygous carrier parents and their two affected children in Family Two. **(I and J)** Relative abundances of LPG **(I)** and BMP **(J)** species in peripheral blood mononuclear cells (PBMCs) from the family shown in **(F and H)**. In **(E–J)**, the abundance of each lipid species was first normalized to endogenous POPC and then normalized to its mean abundance in the corresponding healthy control group **(E and G)** or carrier-parent group **(F, H, I, and J)**. Gray bars represent healthy controls or carrier parents, and pink bars represent patients or affected children.

## Supplementary Tables

**Table S1:** Untargeted lipidomic analysis of wild-type and *CLN6* knockout HEK293T cells. Related to **Fig. 1A**.

**Table S2:** MRM transitions for targeted analysis of unlabeled BMP, LPG, and PG.

**Table S3:** MRM transitions for targeted analysis of derivatized BMP, LPG, and PG.

## Materials and Methods

### Antibodies

Rabbit anti-Strep (MA-46974, Thermo Fisher Scientific, Waltham, MA), mouse anti-Strep (MA5-17282, Thermo Fisher Scientific), rabbit anti-CLN6 (20315-1-AP, Proteintech, Rosemont, IL), mouse anti-vinculin (SC-73614, Santa Cruz Biotechnology, Dallas, TX), rabbit anti-CLN5 (ab170899, Abcam, Cambridge, UK) were used in western blot at a 1:1000 dilution. Mouse anti-GM2 (A2576, TCI Chemicals, Tokyo, Japan) was used in immunostaining at a 1:200 dilution.

### Plasmids sequence

The following codon optimized CLN6 WT and mutant geneblocks were cloned to lentivectors or transient expressing vector

For lentiviral gene delivery

CLN6-2×Strep WT

ATGGAAGCAACACGCCGTAGACAACACTTAGGGGCAACGGGCGGACCTGGTGCGCAACTTGGGGCTAGTTTCTTGCAAGCACGGCACGGTTCAGTGTCAGCCGACGAGGCTGCACGCACCGCCCCGTTTCATTTAGACCTGTGGTTTTATTTCACACTGCAGAACTGGGTCCTGGACTTTGGAAGACCAATTGCCATGTTAGTGTTCCCTCTGGAGTGGTTTCCGCTTAACAAACCGTCTGTCGGAGATTATTTCCACATGGCCTATAATGTTATTACTCCATTCCTGCTGTTAAAATTGATTGAGCGCTCTCCAAGAACCCTTCCCAGGTCCATAACTTACGTCAGCATTATTATTTTCATCATGGGCGCTTCTATACATTTAGTGGGTGACTCCGTCAATCATAGACTCCTCTTTTCCGGCTACCAGCACCATCTCAGTGTGAGAGAAAACCCAATAATTAAGAACTTGAAACCGGAAACCCTTATTGATTCATTTGAGCTCCTGTATTATTATGATGAATATCTGGGCCACTGCATGTGGTATATCCCCTTCTTCCTGATCTTGTTTATGTATTTCAGCGGATGCTTTACCGCTAGTAAAGCAGAGTCCCTTATCCCTGGACCCGCCTTATTGCTTGTCGCTCCATCAGGACTTTACTATTGGTATCTTGTAACGGAAGGACAAATTTTCATACTGTTTATCTTCACCTTCTTCGCAATGTTAGCCCTCGTCTTACACCAGAAACGCAAACGCTTGTTCTTGGATTCAAATGGATTGTTTCTGTTTTCTTCATTTGCCCTCACGTTGCTGCTGGTTGCCCTTTGGGTGGCCTGGCTGTGGAATGATCCAGTTCTCAGGAAGAAGTATCCCGGGGTTATATACGTACCAGAGCCCTGGGCCTTCTACACCCTGCATGTTTCTAGCAGACACGGTGGAGGAGGCTCTGGGGGGGGTGGATCAGGTGGCTCTTGGTCTCATCCGCAATTTGAGAAAGGCGGCGGCGGATCTGGGGGGGGCGGCTCCGGCGGTAGTGCTTGGTCTCATCCTCAGTTTGAAAAGTAA

CLN6-2×Strep Y182AY223A

ATGGAAGCAACTCGACGAAGACAACACTTGGGCGCAACTGGGGGCCCGGGTGCACAGCTTGGCGCGAGTTTCCTGCAAGCGAGGCATGGCTCCGTGTCCGCTGATGAGGCCGC AAGGACGGCACCTTTCCATTTGGATTTATGGTTTTACTTCACCCTCCAAAATTGGGTATTAGATTTCGGCAGACCCATCGCAATGTTGGTCTTTCCTCTTGAGTGGTTCCCGCTGAATAAGCCATCCGTAGGCGACTATTTCCACATGGCTTACAACGTGATTACACCTTTCCTCCTCCTGAAGCTGATAGAACGGTCCCCTCGGACGCTTCCCAGGTCAATAACTTATGTGAGCATTATCATATTCATTATGGGGGCGAGCATCCATCTTGTCGGCGACTCCGTCAACCACCGCCTGCTCTTTTCAGGCTACCAGCACCATCTTTCCGTCAGAGAAAATCCTATAATAAAAAACCTTAAGCCGGAGACACTGATAGACTCCTTTGAACTCCTGTACTACTATGACGAATACCTGGGACACTGTATGTGGGCTATCCCTTTTTTTCTGATCCTGTTCATGTACTTCAGCGGCTGCTTTACCGCCTCTAAGGCTGAATCACTTATACCTGGGCCAGCTTTGCTGCTTGTGGCACCCAGCGGACTGTACTATTGGGCCCTGGTGACAGAGGGGCAAATATTCATTCTGTTTATCTTTACTTTCTTTGCAATGCTTGCCTTGGTGCTCCACCAGAAACGAAAGCGACTGTTCCTGGACAGTAACGGCCTGTTCCTCTTCTCAAGTTTTGCACTGACGCTGCTCCTGGTTGCTTTATGGGTGGCCTGGCTGTGGAATGACCCAGTCCTGCGGAAGAAGTACCCAGGTGTCATCTACGTGCCAGAACCCTGGGCTTTCTACACTTTACACGTGAGTTCTCGGCATGGGGGAGGCGGGAGCGGTGGCGGAGGATCTGGCGGATCCTGGTCTCATCCCCAGTTCGAAAAAGGAGGAGGCGGATCTGGCGGTGGCGGATCCGGTGGCAGTGCGTGGAGTCATCCGCAGTTTGAAAAATGA

CLN6-2×Strep D173Q/E174Q/H178A

ATGGAGGCTACAAGAAGGCGACAGCACCTCGGTGCAACCGGTGGACCTGGAGCTCAGCTGGGTGCATCATTTCTCCAGGCACGCCACGGAAGCGTGTCCGCTGACGAGGCTGCAAGAACAGCCCCATTCCACCTGGATCTGTGGTTCTACTTCACACTTCAGAATTGGGTTCTGGACTTCGGAAGGCCCATCGCTATGCTGGTTTTCCCCCTGGAGTGGTTCCCCCTGAACAAACCTTCCGTGGGTGATTATTTCCATATGGCCTACAACGTTATCACACCCTTTCTGTTGCTGAAGCTGATTGAGAGATCCCCCCGAACTCTTCCCCGCTCAATCACTTACGTGAGTATCATCATTTTCATCATGGGTGCCAGCATCCATTTGGTCGGCGACAGTGTTAACCACCGGCTGCTGTTCTCAGGCTATCAGCATCACCTCTCTGTGCGCGAGAATCCAATAATCAAAAACCTGAAGCCTGAAACACTGATAGACAGTTTCGAACTGCTGTATTATTACCAGCAATATCTCGGAGCCTGTATGTGGTATATCCCATTCTTCCTGATACTCTTCATGTATTTCAGCGGGTGTTTCACCGCTTCCAAGGCTGAAAGCTTGATCCCCGGACCTGCCCTCCTGCTGGTTGCACCATCTGGACTCTATTATTGGTACTTGGTGACCGAGGGCCAGATCTTCATCCTGTTTATTTTCACCTTCTTCGCCATGTTGGCCCTGGTCCTTCATCAAAAGAGGAAGCGACTGTTTCTCGATAGTAATGGACTGTTCCTTTTTTCATCCTTTGCCCTGACACTGCTCCTGGTTGCGCTCTGGGTCGCGTGGTTGTGGAATGACCCCGTCTTAAGAAAAAAATATCCAGGGGTCATCTACGTGCCTGAGCCCTGGGCATTCTACACACTGCATGTTTCTTCTAGACATGGTGGGGGGGGCTCTGGGGGTGGCGGCTCCGGTGGTAGTTGGAGTCATCCACAGTTCGAGAAAGGAGGAGGTGGAAGCGGTGGAGGGGGATCTGGTGGATCTGCCTGGAGCCACCCTCAATTCGAAAAATGA

CLN6-2×Strep D173A/E174A/H178A

ATGGAAGCCACACGAAGGCGCCAACATCTGGGCGCTACGGGGGGGCCCGGTGCACAACTGGGAGCTAGTTTCCTGCAGGCTAGGCACGGATCTGTGAGTGCAGATGAGGCCGCCAGGACCGCCCCCTTCCATCTTGACCTGTGGTTCTACTTTACTTTACAGAACTGGGTCTTAGACTTCGGAAGGCCCATTGCAATGTTAGTCTTTCCTCTGGAGTGGTTCCCGCTGA ATAAGCCTAGCGTGGGCGATTACTTTCACATGGCCTACAATGTAATCACTCCTTTCCTGTTACTGAAACTCATCGAGCGCTCTCCCCGCACACTGCCAAGGAGCATCACTTACGTTTCCATCATCATCTTCATAATGGGCGCTTCCATACACCTCGTGGGGGATTCCGTGAATCATCGACTCTTGTTTTCCGGTTACCAGCATCACCTGTCCGTGCGGGAAAATCCTATCATCAAAAACCTCAAACCCGAAACGCTCATCGACAGTTTCGAATTACTGTATTATTACGCTGCGTATCTGGGAGCCTGCATGTGGTACATTCCTTTCTTCTTGATTCTGTTCATGTACTTCAGTGGCTGCTTCACCGCCTCCAAGGCAGAAAGTCTGATCCCAGGACCAGCTCTGCTCTTGGTTGCACCAAGTGGGCTGTATTATTGGTACCTTGTTACCGAGGGACAGATCTTCATTCTCTTTATTTTTACCTTCTTCGCCATGCTCGCGTTAGTGCTGCACCAAAAAAGGAAAAGATTGTTTTTAGATTCAAACGGATTGTTTCTCTTCAGCAGTTTCGCTCTTACTCTGCTGTTGGTCGCCTTGTGGGTCGCGTGGCTGTGGAACGATCCAGTGCTTCGTAAAAAATACCCAGGAGTCATTTACGTTCCAGAGCCCTGGGCCTTCTATACACTGCATGTGAGCTCTAGGCACGGGGGAGGGGGATCCGGCGGGGGGGGCTCAGGCGGTAGCTGGTCACACCCTCAATTTGAAAAGGGTGGCGGCGGATCCGGTGGAGGCGGTAGCGGCGGAAGCGCTTGGAGCCACCCACAGTTTGAAAAGTAA

CLN6-2×Strep R62H

ATGGAAGCGACTCGACGTCGCCAGCACCTCGGCGCGACTGGAGGTCCCGGCGCCCAGCTGGGCGCATCTTTCTTGCAGGCAAGACATGGGTCTGTCTCTGCTGATGAGGCTGCCCGCACTGCTCCCTTTCATTTGGATCTGTGGTTCTACTTCACTCTCCAAAATTGGGTGCTGGATTTTGGACATCCCATCGCCATGCTTGTGTTCCCCCTGGAGTGGTTTCCGTTGAATAAGCCCTCTGTCGGCGATTACTTTCATATGGCTTACAATGTCATCACACCATTTCTTCTGCTCAAGCTTATAGAGCGTAGCCCGAGGACTTTGCCCCGGTCTATAACATATGTATCCATTATCATCTTTATTATGGGAGCGAGTATACATCTCGTCGGCGACTCCGTTAACCATAGGCTGCTTTTTTCCGGATATCAACACCATCTGAGCGTGCGGGAGAACCCAATTATCAAGAATCTTAAGCCCGAGACGTTGATTGACAGCTTTGAGCTGCTCTACTATTATGATGAATACCTCGGGCACTGTATGTGGTACATACCCTTTTTCCTCATCCTTTTTATGTATTTTTCCGGTTGTTTCACTGCATCTAAGGCGGAATCTTTGATTCCCGGGCCGGCACTTTTGCTCGTCGCACCGAGTGGCCTGTACTATTGGTATCTCGTCACCGAGGGTCAGATCTTTATTCTCTTCATCTTCACATTTTTCGCAATGCTGGCACTTGTTCTCCATCAGAAGAGAAAGCGTCTGTTCCTCGATTCTAATGGTCTCTTCCTTTTCTCTAGCTTTGCCCTGACCTTGCTTCTGGTGGCTCTTTGGGTGGCCTGGCTCTGGAACGACCCAGTCCTGAGGAAGAAATATCCTGGAGTCATATACGTCCCTGAACCATGGGCCTTTTACACCCTTCACGTCTCTTCTCGTCACGGCGGCGGTGGTAGCGGTGGGGGCGGTTCTGGCGGATCCTGGTCCCACCCCCAGTTTGAGAAGGGAGGCGGTGGGAGCGGCGGGGGTGGCTCAGGCGGGTCAGCCTGGAGTCATCCTCAGTTTGAGAAATGA

CLN6-2×Strep H127Y

ATGGAAGCTACCAGACGCAGACAGCATCTGGGCGCAACCGGTGGTCCAGGAGCCCAGCTGGGTGCGTCATTCTTGCAAGCCCGCCACGGTAGTGTGTCTGCAGATGAGGCAGCGCGAACGGCCCCCTTCCACCTCGACCTGTGGTTTTACTTCACCCTCCAAAATTGGGTCCTCGACTTCGGACGGCCAATCGCCATGCTCGTTTTCCCATTAGAGTGGTTCCCCCTGAATAAACCGAGCGTGGGAGATTATTTCCACATGGCTTATAACGTGATCACTCCATTTCTTCTGCTGAAACTCATCGAGAGGAGCCCTCGAACCCTGCCCCGGTCAATAACTTACGTG AGCATCATCATTTTCATTATGGGTGCTAGCATTTATCTCGTTGGGGACTCAGTCAACCATCGGCTTCTGTTTAGTGGCTACCAGCATCACCTGTCCGTGCGAGAGAATCCCATCATCAAGAACCTGAAACCGGAGACACTGATTGATAGCTTCGAGCTTCTCTACTACTACGACGAGTATTTGGGCCATTGTATGTGGTACATACCCTTCTTCTTAATCCTGTTCATGTACTTCAGTGGATGCTTTACCGCTAGTAAAGCTGAGAGCCTGATCCCTGGGCCAGCTCTCCTGCTGGTCGCTCCAAGCGGCCTTTACTATTGGTACTTGGTCACAGAAGGACAAATTTTTATTTTGTTCATTTTCACCTTTTTCGCTATGTTAGCACTCGTCCTGCATCAGAAACGAAAGCGATTATTCCTCGATTCCAACGGGCTCTTCTTGTTCAGCTCTTTTGCCCTTACCCTTCTCTTGGTCGCTCTCTGGGTCGCCTGGCTTTGGAACGACCCAGTGTTGCGCAAGAAGTATCCCGGCGTCATTTATGTCCCTGAGCCCTGGGCTTTTTACACCTTGCATGTTTCCAGCCGTCACGGAGGGGGCGGATCCGGAGGAGGCGGCAGCGGAGGATCTTGGAGCCACCCACAGTTTGAAAAGGGTGGTGGTGGCTCCGGTGGAGGTGGTTCAGGGGGGAGCGCCTGGAGTCATCCACAGTTCGAAAAGTAA

For protein purification

CLN6-2×Strep WT

ATGGAAGCAACCCGTAGAAGGCAGCACCTGGGCGCAACTGGGGGACCAGGCGCTCAGCTGGGAGCTTCATTCCTGCAAGCGCGACATGGGTCAGTGTCTGCTGATGAAGCGGCAAGAACCGCCCCGTTTCACCTGGACCTCTGGTTCTATTTCACCCTCCAGAATTGGGTCTTAGACTTCGGGCGCCCTATTGCCATGTTGGTCTTTCCACTGGAGTGGTTTCCCCTCAATAAACCAAGCGTCGGAGACTACTTCCACATGGCTTACAACGTGATCACTCCTTTCCTCCTTCTGAAGCTCATCGAGCGCTCACCAAGGACTCTGCCGAGAAGCATTACCTATGTGTCCATCATTATCTTTATAATGGGGGCGTCAATCCATCTGGTCGGCGATAGCGTGAACCACAGACTCCTCTTTTCCGGCTATCAGCACCACCTCAGCGTCCGGGAGAACCCCATAATCAAAAATTTGAAGCCTGAGACGCTCATCGACTCTTTCGAACTCCTCTACTACTACGATGAATATCTCGGACATTGCATGTGGTATATCCCATTCTTCCTGATTCTGTTTATGTACTTCAGCGGCTGCTTCACAGCGTCCAAGGCCGAGTCACTGATTCCTGGACCCGCACTGCTTTTAGTCGCGCCGTCTGGGCTGTACTACTGGTATCTCGTCACCGAAGGCCAAATATTCATCCTGTTTATTTTCACATTTTTCGCTATGCTGGCCTTAGTGCTCCACCAAAAAAGGAAGCGACTTTTCCTGGACTCAAACGGGCTGTTCCTCTTTTCAAGCTTTGCGCTCACACTGCTGCTGGTGGCCCTGTGGGTGGCCTGGCTTTGGAATGACCCAGTCCTCCGGAAGAAATATCCCGGTGTGATCTACGTGCCTGAGCCTTGGGCCTTCTATACGCTTCACGTATCTTCACGCCACGGGGGAGGTGGGAGCGGAGGCGGGGGGAGCGGCGGGTCATGGTCACATCCTCAGTTTGAGAAAGGCGGTGGCGGATCTGGGGGGGGAGGGTCCGGAGGCTCCGCCTGGAGCCACCCTCAGTTTGAGAAGTAG

CLN6-2×Strep H127Y

ATGGAGGCGACCCGCCGTCGGCAGCATTTAGGAGCCACCGGCGGCCCAGGCGCCCAACTGGGCGCCTCTTTTTTGCAGGCAAGACACGGAAGCGTTTCCGCAGACGAAGCAGCGCGAACGGCTCCTTTTCACCTTGACTTGTGGTTTTATTTTACCCTGCAGAACTGGGT TCTCGACTTCGGCAGACCGATCGCCATGCTGGTTTTTCCTTTAGAGTGGTTCCCACTCAATAAACCGTCCGTCGGTGACTATTTCCACATGGCCTACAACGTAATTACCCCCTTCCTGCTGCTTAAACTTATCGAAAGATCACCAAGAACGCTGCCAAGGTCCATAACTTATGTTTCAATCATTATTTTCATTATGGGAGCAAGCATTTACCTGGTAGGGGATAGCGTCAACCACAGGCTTCTCTTCAGCGGCTATCAGCACCATCTGAGCGTTCGCGAAAACCCCATCATTAAAAACTTGAAACCCGAGACTCTCATCGATTCTTTCGAACTCCTCTATTACTACGATGAATATCTGGGCCATTGCATGTGGTATATACCATTTTTTCTCATCCTTTTCATGTATTTTAGCGGATGTTTCACAGCATCTAAGGCTGAGTCCCTCATTCCTGGGCCCGCGCTTCTGCTTGTTGCCCCATCCGGGCTCTACTACTGGTACCTGGTCACCGAGGGTCAGATCTTTATTCTGTTTATCTTTACTTTCTTTGCCATGCTTGCTTTGGTACTCCACCAGAAACGCAAACGTCTGTTTCTGGATAGCAACGGCCTTTTTCTCTTCTCTTCATTTGCCCTGACTCTCCTTTTAGTGGCCTTGTGGGTGGCTTGGCTCTGGAATGATCCTGTGTTGCGGAAGAAGTACCCGGGTGTCATCTATGTGCCGGAGCCCTGGGCATTCTATACCCTTCATGTGTCTTCCAGGCACGGAGGCGGGGGGTCTGGTGGCGGGGGGAGCGGGGGGTCTTGGAGCCATCCTCAGTTCGAAAAAGGCGGAGGGGGAAGCGGAGGGGGCGGATCCGGGGGGTCCGCATGGTCCCACCCACAATTCGAAAAGTAA

### Animal studies

*Cln6^nclf^* mice (*35*) (The Jackson Laboratory, Strain: #003605) were maintained on a C57B6/J background in the AAALAC-accredited rodent facility at Massachusetts General Hospital (MGH). Heterozygous matings were used to generate homozygous experimental animals and control littermates (heterozygous), which were aged to 2.9 months for euthanasia and tissue collection. Mice were euthanized by inhalation of CO_2_-bottled gas, in accordance with current guidelines from the American Veterinary Medical Association. Isolated tissues were snap frozen and stored at –80°C until further processing. All mouse procedures used for this study were reviewed and approved by the Subcommittee on Research Animal Care at MGH to ensure compliance with government regulations and animal welfare guidelines (Protocol # 2008N000013).

### Cell culture

HEK293T cells (American Type Culture Collection CRL-3216) and their derivatives were cultured in Dulbecco’s modified Eagle’s medium (DMEM) (11965092, Thermo Fisher Scientific) supplemented with 10% heat-inactivated fetal bovine serum (FBS) (16140071, Thermo Fisher Scientific), 100□IU/ml penicillin and 100□µg/ml streptomycin (SV30010, Cytiva, Marlborough, MA). ARPE-19/HPV-16 cells (American Type Culture Collection CRL-2502) and their derivatives were cultured in 1:1 mixture of Dulbecco’s Modified Eagle Medium and Ham’s F-12 medium (DMEM:F12 1:1) (11330057, Thermo Fisher Scientific). HEK293T and ARPE19/HPV-16 were both maintained at 37□°C in an incubator with 5% CO_2_. For protein expression, Expi293F cells (Thermo Fisher Scientific) were maintained in a shaking incubator at 37□°C with 8% CO_2_. These cells were routinely tested for mycoplasma contamination.

### Generation of knockout cell lines using CRISPR-Cas9 technology

*CLN6* knockout HEK293T and ARPE-19/HPV16 cells were generated following the Editco Gene Knockout Kit protocol using the following mixed guide sgRNA sequences: UGGGGUUCUCACGGACAGAC, GUCACCCACCAGGUGGAUGC and GUACGUGAUGGAGCGUGGCA. *CLN8* knockout were generated with sgRNA CAAUGAAUCCUGCGAGCGAU, CGCUCCACGCUGAUGGUCGC and AGAACGGUAAGUGGCAUUCA. In short, a combined 180-pmol sgRNA and 20-pmol Cas9 (Integrated DNA Technologies) was incubated at room temperature for 10 min. Concurrently, HEK293Ts were washed with PBS, detached using enzyme-free dissociation buffer (Thermo Fisher Scientific, 13151014), and counted to determine their density. A suspension of 150,000 cells was added to the newly formed ribonucleoprotein complexes, transferred to a Lonza Nucleocuvette, and electroporated using the CM-130 (HEK293T) and EA-104 (ARPE-19/HPV-16) program code. Following electroporation, the cells were placed in growth medium and transferred to a 24-well plate. Single-cell populations were obtained using fluorescence-activated cell sorting by FACSAria II (BD, Franklin Lakes, NJ), and the sgRNA target sites were sequenced after PCR amplification. Primers TTGGTGAAAGGACAGGTGCG and CCTGCTAAAGGGACCAGCC were used to amplify CLN6 targeted genome. Primers TGTGTCCTTGTTTCTATCGAGTCA and CGTGATGTGAAACCAGCACC were used to amplify CLN8 targeted genome. Finally, genetic knockouts were confirmed using Synthego ICE CRISPR analysis, an insertion and deletion deconvolution web-based tool. For CLN6 knockout, we also used western blot to confirm protein deletion.

### Expression and purification of proteins

The coding sequences of human CLN6 or its mutants were cloned into the pTwist-CMV vector (Twist Bioscience, South San Francisco, CA) in frame with a C-terminal Twin-Strep-tag (2×Strep). All constructs were verified by whole-plasmid sequencing. Expi293F cells (Thermo Fisher Scientific, Waltham, MA) were grown in Expi293 Expression Medium at 37 °C and 8% CO_2_ with orbital shaking at 125 rpm and were transfected at a density of 3 × 10^6 cells/ml with 0.5 µg of plasmid DNA per ml of culture using PEI STAR (Tocris Bioscience, Bristol, UK) at a 5:1 (w/w) PEI:DNA ratio. Cells were harvested 72 hours after transfection by centrifugation at 500 × g for 10 min at 4 °C and cell pellets were resuspended in lysis buffer [50 mM HEPES pH 7.4, 150 mM NaCl, 5% (v/v) glycerol, 1% (w/v) lauryl maltose neopentyl glycol (LMNG; Anatrace, Maumee, OH), 0.5% (w/v) cholesteryl hemisuccinate (CHS; Anatrace), and cOmplete EDTA-free protease inhibitor cocktail (Roche, Basel, Switzerland)] and solubilized for 2 hours at 4 °C with rotation. The lysate was centrifuged at 135,000 × g for 1 hour at 4 °C in a Ti-32 rotor (Beckman Coulter).

The clarified supernatant was then collected and passed through a 5-ml Strep-Tactin XT 4Flow high-capacity column (IBA Lifesciences, Göttingen, Germany) pre-equilibrated in wash buffer [100 mM Tris-HCl pH 8.0, 150 mM NaCl, 1 mM EDTA, 0.1% (w/v) LMNG, 0.05% (w/v) CHS] at 4 °C. The column was washed with 10 × column volumes (CV) of wash buffer, followed by a 10 × CV of wash buffer supplemented with 10 mM ATP and 20 mM MgCl_2_ to displace co-purifying chaperones, and then with 3 × CV of wash buffer to remove residual nucleotide. Bound protein was eluted with 3 × CV of elution buffer [100 mM Tris-HCl pH 8.0, 150 mM NaCl, 1 mM EDTA, 50 mM desthiobiotin, 0.1% (w/v) LMNG, 0.05% (w/v) CHS]. Eluted fractions were pooled and concentrated in an Amicon Ultra centrifugal filter unit (3,000 Da molecular weight cutoff) (UFC900324, MilliporeSigma Burlington, MA).

The concentrated eluate was applied on a Superdex 200 Increase 10/300 GL column (Cytiva) equilibrated in size exclusion chromatography (SEC) buffer [50 mM HEPES pH 7.4, 150 mM NaCl, 5% (v/v) glycerol, 0.01% (w/v) LMNG, 0.005% (w/v) CHS] at a flow rate of 0.375 ml/min at 4 °C. 0.5-ml fractions were collected for 23 ml. Fraction purity was assessed by SDS-PAGE with Coomassie blue staining. Fractions containing pure CLN6-2×Strep were pooled and concentrated as above; protein concentration was determined from the absorbance at 280 nm on a NanoDrop 2000 spectrophotometer (Thermo Fisher Scientific) using a calculated extinction coefficient of 86080 M^-1^cm^-1^, with SEC buffer as the blank. Aliquots of 20 µl were added into 0.2-ml tubes, flash-frozen in liquid nitrogen, and stored at –80 °C.

### Lentivirus preparation and infection

WT and mutant CLN6 coding sequences were cloned into pTwist-Lenti-CMV-Puro (Twist Bioscience, South San Francisco, CA) and verified by whole-plasmid sequencing. Lentiviral particles were produced in HEK293T cells. Cells at 70–80% confluence in a 10-cm dish were co-transfected with 6 µg of the lentiviral vector, 5 µg of the packaging plasmid psPAX2 (Addgene plasmid #12260) and 1 µg of the VSV-G envelope plasmid pMD2.G (Addgene plasmid #12259) using X-tremeGENE HP DNA transfection reagent (MilliporeSigma, 06365787001) at a 3:1 (v/w) reagent:DNA ratio according to the manufacturer’s instructions. 18 hours after transfection, the medium was replaced with DMEM containing 30% (v/v) FBS, and virus-containing supernatant was harvested 48 hours after transfection. The supernatant was clarified by centrifugation at 500 × g for 10 min at 4 °C followed by filtration through a 0.45-µm filter and concentrated with Lenti-X Concentrator (Takara Bio, Shiga, Japan, 631232) by adding one-third volume of reagent, incubating overnight at 4 °C, and centrifuging at 1,500 × g for 45 min at 4 °C. Viral pellets were resuspended in 1/20 of the original volume of complete DMEM and used immediately or stored in single-use aliquots at −80 °C. Target cells were seeded at 0.5 million cells per well of a 6-well plate and transduced during seeding with 100 µL concentrated virus in the presence of 10 µg/ml polybrene (MilliporeSigma, TR-1003-G). The virus-containing medium was removed 24 hours after transduction and replaced with fresh growth medium. 72 hours after transduction, transduced cells were selected in medium containing 2 µg/ml puromycin for 2 days, with non-transduced cells processed in parallel as a control. Stable polyclonal populations were expanded, and CLN6 expression was confirmed by western blot.

### Stem cell maintenance and differentiation into iNeurons

Human induced pluripotent stem cells (iPSCs; KOLF2.1J) were obtained from The Jackson Laboratory. iPSCs were maintained on Matrigel-coated plates (Corning, 354230, Corning, NY) in mTeSR Plus medium (STEMCELL Technologies, 100-0276, Vancouver, BC, Canada) supplemented with 50 U/mL penicillin-streptomycin (Thermo Fisher Scientific, 15140122). Cells were fed every other day and passaged every 3–4 days using ReLeSR (STEMCELL Technologies, 100-0483) according to the manufacturer’s instructions. Following passaging, cells were cultured in mTeSR Plus medium supplemented with 10 µM ROCK inhibitor (Tocris, 1254, Bristol, UK). Within 24 hours, the medium was replaced with fresh medium lacking the ROCK inhibitor.

iPSCs transduced with a Tet-On inducible system to drive expression of the transcription factor NGN2 were induced in DMEM/F12 medium (Thermo Fisher Scientific, 11320033) supplemented with 1 × N2 (Thermo Fisher Scientific, 17502-048) and 1 µg/mL doxycycline (MilliporeSigma, D5207). On day 3 of differentiation, cells were dissociated and replated onto tissue culture plates coated with poly-D-lysine and laminin. Cells were maintained in Neurobasal medium (Thermo Fisher Scientific, 21103049) supplemented with 1 × B-27 (Thermo Fisher Scientific, 17504044) and the neurotrophic factors brain-derived neurotrophic factor (BDNF; PeproTech, 450-10, Cranbury, NJ) and glial cell line-derived neurotrophic factor (GDNF; PeproTech, 450-03). Half-media changes were performed every 3–4 days.

### Deuterated GPG tracing

Synthetic d5-*R,S*-GPG and d5-*S,S*-GPG were dissolved in ultrapure water to prepare a 40 mM stock solution.

For tracing experiments in HEK293T cells, 0.5 million cells were seeded per well in poly-L-lysine-coated 6-well plates. The following day, the desired amount of GPG or an equal volume of water was added to the culture medium. After 48 hours of treatment, cells were harvested for metabolite or lipid extraction.

For supplementation experiments in iNeurons, 1 million iPSCs were seeded in 6-well plates and differentiated according to the protocol described above. On day 10 after differentiation induction, cells were treated with increasing concentrations of *S,S*-GPG up to 300 µM or an equivalent volume of water as the vehicle control for 48 hours. The iNeurons were then harvested for lipid extraction.

For tracing experiments in lysate, HEK293T cells of the indicated genotypes were grown to 80–90% confluence in 15 cm dishes, washed once with ice-cold PBS, and collected. Cells were resuspended in ultrapure water (Thermo Fisher Scientific, 10977023) and homogenized with 25 strokes of a Dounce homogenizer (VWR, 89026, Radnor, PA). Homogenates were centrifuged at 1,000 × g for 2 min at 4 °C, and the supernatant was collected as the lysate. Each reaction contained 37 µL of lysate, 100 µM d5-*R,S*-GPG tracer, and 500 µM CaCl, with or without 2 mM BAPTA (Ambeed, Buffalo Grove, IL), in 1× PBS (diluted from 10× PBS) in a final volume of 50 µL. Reactions were incubated overnight at 37 °C.

### Human subjects and patient-derived samples

All studies involving human material from Stanford were approved by the Institutional Review Board of Stanford University (protocol no. 34131, a.k.a CGID/BMT328). All studies involving human material from Hamburg, Germany were approved by the local medical ethics committee of the Ärztekammer Hamburg (PV7199). Written informed consent was obtained from all participants or, for minors, from their parents or legal guardians, including consent for publication of anonymized genetic and biochemical data. Samples were obtained from three unrelated families affected by CLN6-associated neuronal ceroid lipofuscinosis. Family One comprised two affected individuals carrying the homozygous in-frame deletion NM_017882.3(CLN6):c.461_463del [p.(Ile154del)], which removes a single isoleucine residue of CLN6 and has been reported in individuals with late-infantile CLN6 disease. Family Two comprised two affected individuals carrying homozygous NM_017882.3(CLN6):c.794_796del [p.(Ser265del)]. Family Three comprised one affected individual carrying NM_017882.3(CLN6):c.542+5G>T (previously designated IVS5+5G>T). Dry blood spots were obtained from Family One and Two. Peripheral blood mononuclear cells (PBMCs) were isolated from Family Two. For each affected individual and parent, 2 ml of EDTA-anticoagulated whole blood was processed by density-gradient centrifugation over Ficoll-Paque PLUS (Cytiva) within 6 hours of venipuncture, washed twice in PBS (pH 7.4), and cells were then counted and pelleted, pellets were flash frozen with liquid nitrogen before lipid extraction. Fibroblasts were obtained from Family Three and three healthy controls.

### Immunoblotting

Cells were washed once with ice-cold phosphate-buffered saline and lysed on ice in lysis buffer (50 mM HEPES pH 7.4, 40 mM NaCl, 2 mM EDTA, 1% (v/v) Triton X-100, 1.5 mM sodium orthovanadate, 30 mM sodium fluoride, 10 mM sodium β-glycerophosphate) supplemented with cOmplete EDTA-free protease inhibitor cocktail (Roche, Basel, Switzerland) for 20 min. Lysates were cleared by centrifugation at maximum speed for 10 min at 4 °C. The supernatant of lysates were mixed with 5× SDS sample buffer containing 10% β-mercaptoethanol, incubated at 37 °C for 5 min and resolved on 4–20% Bis-Tris SDS-PAGE gels (Thermo Fisher Scientific) at 120 V for ∼90 min. Proteins were transferred to 0.22-µm nitrocellulose membranes (LI-COR Biosciences, Lincoln, NE) by wet transfer at 60 V for 2 hours in transfer buffer (25 mM Tris, 192 mM glycine, 20% (v/v) methanol). Membranes were blocked in 5% non-fat milk in Tris-buffered saline containing 0.1% (v/v) Tween 20 (TBST) for 1 hour at RT, then incubated overnight at RT with primary antibodies diluted in TBST containing 5% non-fat milk. Membranes were washed three times for 5 min per wash with TBST and incubated for 1 hour at room temperature with the appropriate IRDye 680RD– and IRDye 800CW-conjugated secondary antibodies (LI-COR Biosciences) diluted 1:10,000. After three additional 5-min TBST washes, membranes were imaged on an Odyssey CLx near-infrared imaging system (LI-COR Biosciences), and pictures were acquired with Image Studio software (LI-COR Biosciences).

### Fluorescent microscopy and analysis

One million WT or *CLN6*-knockout ARPE-19/HPV-16 cells were seeded in 10-cm dishes and cultured for 8 days in complete DMEM/F-12 supplemented with either H_2_O or 400 µM *S,S*-GPG. Cells were then reseeded at 20,000 cells per well in eight-chamber cell culture slides (CELLTREAT), with each well containing 500 µL of complete DMEM/F-12. The cells were continued to be treated with H_2_O or 400 µM *S,S*-GPG for an additional 2 days. Cells were subsequently fixed with 300 µL of 4% paraformaldehyde for 15 min and washed three times with 600 µL of PBS. For filipin staining, cells were incubated with 200 µL of 1 mg/mL filipin (Cayman Chemical). For GM2 immunostaining, cells were first blocked with 5% nonfat milk in PBS and then incubated overnight at 4°C with an anti-GM2 antibody diluted 1:200. The following day, cells were washed three times with PBS and incubated for 1 hour with Alexa Fluor 488-conjugated goat anti-mouse secondary antibody (#4408, Cell Signaling Technology) diluted 1:1,000. Finally, the slides were mounted with mounting medium and glass coverslips and subjected to fluorescence imaging.

Images were acquired on a Zeiss LSM 980 confocal microscope (ZEN 3.6) with a 25X oil immersion objective. Alexa Fluor 488 was excited with a 488 nm laser line and detected at 517 nm emission; Filipin was excited with a 405 nm laser line and detected at 465 nm emission. Images were captured at 1024 x 1024 pixel resolution (pixel size 0.331 µm) in 8-bit depth using a multi-channel photomultiplier tube detector (MA-Pmt1). Laser power and detector gain were kept constant across all conditions to ensure comparability of fluorescence intensity measurements. Cell segmentation and fluorescence quantification were performed using scikit-image (v0.25.2) and SciPy (v1.15.0). Raw.czi files were read directly without preprocessing or compression; all original pixel values were preserved. For publication archiving, images were converted to TIFF format (lossless, no compression) with verification of pixel-level integrity.

### Schrodinger Docking

The structure of CLN6 (Uniprot: Q9NWW5) coordinated with a calcium ion was predicted using AlphaFold 3 (*43*). The predicted structure was loaded into Maestro software (Schrödinger Release 2026-2). Protein was prepared for docking using the protein preparation wizard to add hydrogens, create disulfide bonds, and assign bond orders. Because the model was generated computationally, no crystallographic waters were present, and no side chains were missing. Ionization and tautomeric states were generated at pH 7.4 ± 0.5 to reflect the ER lumen pH. The structure was subjected to restrained minimization with the OPLS4 force field to an RMSD of 0.30 Å. The receptor grid was generated centroid of the selected residues Arg 62 and His 127 corresponding to known patient mutations in the central cavity of CLN6, this was generated without the ligand and was set to dock ligands with a length of 20 Å. A 2D GPG structure was drawn in ChemDraw, and LigPrep was used to generate possible states at target pH 7.4 ± 0.5 via Epik, tautomers, and stereoisomers in an OPLS4 force field (*44*). For docking the GPG library into CLN6, the standard precision (SP) docking mode of Glide (*45*) was used under the following parameters: flexible ligand sampling, add Epik (*46*) state penalties to docking score, default van der Waals radii scaling, and perform post-docking minimization, retaining 15 poses per ligand for minimization and writing 5 poses per ligand. Docked poses were displayed using UCSF ChimeraX (*47*).

### Extraction of lipids from all sample types for lipidomic analysis

Lipids were extracted following a previous protocol (*31*). Lipids were extracted using chloroform/methanol (2:1, v/v) containing SPLASH LIPIDOMIX isotopically labeled internal standards (Avanti) diluted 1:1,000 (hereafter referred to as lipid extraction buffer).

Cells were harvested in 200 µL of cold methanol/water (4:1, v/v), and the suspension was added to 1,000 µL of lipid extraction buffer. Samples were vortexed at 4 °C for 1 h, followed by the addition of 200 µL of 0.9% NaCl solution and further vortexing at 4 °C for 10 min. Samples were then centrifuged at 3,000 × *g* for 5 min at 4 °C, and 600 µL of the lower chloroform phase was collected. Extracts were dried at room temperature in a SpeedVac and stored at −80 °C until analysis.

Mouse brain and lung tissues were homogenized in 400 µL of PBS using a metal bead and a TissueLyser (Qiagen) at 25 Hz for 30 s and 1 min, respectively. A 25-µL aliquot of each homogenate was added to 1,000 µL of lipid extraction buffer and processed as described above.

For dried blood spots (DBS), one spot per patient was excised using scissors and immersed in 1,000 µL of lipid extraction buffer. Samples were processed as described for cells, except that the filter paper was removed before centrifugation.

Pelleted PBMCs and fibroblasts were resuspended in 200 µL of cold methanol/water (4:1, v/v), added to 1,000 µL of lipid extraction buffer, and processed as described for cells.

On the day of lipidomics analysis, dried extracts were reconstituted in 50 µL of acetonitrile/isopropanol/water (13:6:1, v/v/v), vortexed at 4 °C for 10 min, and centrifuged at maximum speed for 15 min at 4 °C. Approximately 40 µL of the supernatant was transferred to an MS vial for analysis.

### Untargeted lipidomics

Lipids were profiled by LC-MS using a Vanquish Horizon UHPLC and the ID-X Orbitrap Tribrid mass spectrometer (Thermo Fisher Scientific). Samples were loaded and separated using an Ascentis Express C18 column (150 × 2.1 mm, 2.7 µm particle size; Merck) coupled to a guard column (5 × 2.1 mm, 2.7 µm particle size; Merck) held at 55 °C. Each lipid extract was analyzed in two separate injections of 4 µL each, one in positive and one in negative ionization mode.

Mobile phase A consisted of 10 mM ammonium formate and 0.1% (v/v) formic acid dissolved in H_2_O/ACN (3:2, v/v). Mobile phase B consisted of 10 mM ammonium formate and 0.1% (v/v) formic acid dissolved in IPA/ACN (9:1, v/v). The flow rate was maintained at 0.26 mL/min. The gradient was as follows: 0–1.5 min, 32–45% B; 1.5–4 min, 45–52% B; 4–5 min, 52–58% B; 5–8 min, 58–66% B; 8–11 min, 66–70% B; 11–14 min, 70–75% B; 14–18 min, 75–97% B; and 18–23 min, 97% B. Mobile phase B was then returned to 32% over 23–24 min and held at 32% until 27 min for column re-equilibration. An additional 3-min equilibration at 32% B preceded each run.

Mass spectrometric analysis was performed using heated electrospray ionization (H-ESI) in positive ion mode with a spray voltage of 3250 V and negative mode with a spray voltage of 3000 V. Sheath, auxiliary, and sweep gas settings were 40, 10, and 1 arbitrary units, respectively. The ion transfer tube and vaporizer temperatures were set to 300 °C and 375 °C, respectively. Full-scan MS spectra were acquired over an *m/z* range of 250–1500 at an Orbitrap resolution of 120,000, with an automatic gain control (AGC) target of 4 × 10 and a maximum injection time of 50 ms. The RF lens was set to 40%, and EASY-IC internal mass calibration was enabled.

Data-dependent MS/MS acquisition used a cycle time of 0.6 s and a precursor intensity threshold of 2.5 × 10^4^. Precursors were isolated using a quadrupole isolation window of 1.5 *m/z* and fragmented by higher-energy collisional dissociation (HCD) at stepped normalized collision energies of 20%, 30%, and 40%. MS/MS spectra were acquired at an Orbitrap resolution of 30,000, with a first mass of *m/z* 75, an AGC target of 5 × 10, and a maximum injection time of 54 ms. Dynamic exclusion was applied for 2.5 s after one precursor selection, with a mass tolerance of ±10 ppm. Isotopic precursors and precursors already selected within the same cycle were excluded. MS and MS/MS spectra were acquired in profile mode.

Unbiased lipid annotation was performed using LipidSearch 5.1 (LS), utilizing MS1 and MS2-dependent annotation of lipids. Lipids were further quantified using Skyline-daily (26.1.1.209) based on identification results from LS. Features were considered significantly enriched or depleted in *CLN6* knockout cells relative to WT cells if the knockout-to-WT abundance ratio was >2 or <0.5, respectively, with *P* < 0.05. BMP annotations required MS/MS spectra of [M+NH_4_] precursor ions dominated by monoacylglycerol (MAG) fragment ions. Species lacking [M+NH_4_] MS/MS spectra but supported by [M−H]□ MS/MS spectra and eluting earlier than the corresponding PG isomers were annotated as putative BMPs.

Identified lipid species, and their quantitative results, fold change, and *P*-values are listed in Table S1.

### Targeted lipidomics

2 µL of lipid samples were analyzed using an Agilent 1290 UHPLC system coupled with an Agilent 6495D triple quadrupole platform (Agilent, Santa Clara, CA, USA). Samples were loaded and separated using a ZORBAX RRHD Eclipse Plus C18 column (100 mm × 2.1 mm, 1.8 µm particle size; Agilent) coupled to a guard column (5 mm × 2.1 mm, 1.8 µm particle size; Agilent) held at 45 °C. Mobile phase A consisted of water/acetonitrile/isopropanol (5:3:2, v/v/v) containing 10 mM ammonium formate and 5 µM medronic acid. Mobile phase B consisted of water/acetonitrile/isopropanol (1:9:90, v/v/v) containing 10 mM ammonium formate. The flow rate was maintained at 0.26 mL/min. The gradient was as follows: 0–5 min, 15–30% B; 5–8 min, 30–50% B; 8–18 min, 50–60% B; 18–23 min, 60–70% B; 23–24 min, 70–100% B; and 24–26 min, 100% B. Mobile phase B was then returned to 15% over 26–26.5 min and held at 15% until 30 min for column re-equilibration.

MS analysis was performed in multiple reaction monitoring (MRM) mode. Source parameters were set as follows: drying gas temperature, 150 °C; drying gas flow rate, 17 L/min; nebulizer pressure, 20 psi; sheath gas temperature, 200 °C; and sheath gas flow rate, 10 L/min. Capillary voltages were 3500 V and 3000 V, and nozzle voltages were 1000 V and 1500 V, for positive and negative ionization modes, respectively. The precursor-to-product ion transitions (*m/z*) used for multiple reaction monitoring (MRM) are listed in **Table S2**.

Peak integration was done using Skyline-daily (26.1.1.209), and peak areas were normalized to the endogenous POPC peak area in each sample.

### Sample preparation for chemical derivatization

Cells, lysates, and protein incubation samples were extracted in a final volume of 250 µL of cold MeOH:H_2_O (4:1, v/v) solution, vortexed for 10 min, and centrifuged for 5 min at maximum speed at 4 °C. The supernatant was collected and evaporated to dryness using a SpeedVac.

Mouse brain and lung tissues were homogenized in 400 µL of PBS using a metal bead and a TissueLyser (Qiagen) at 25 Hz for 30 s and 1 min, respectively. A 40-µL aliquot of each homogenate was added to 200 µL of cold MeOH, vortexed for 10 min, and centrifuged for 5 min at maximum speed at 4□°C. The supernatant was collected and evaporated to dryness using a SpeedVac.

For DBS, one spot per patient was excised using scissors and immersed in 1000 µL cold MeOH:H_2_O (4:1, v/v) solution. Samples were processed as described for cells, except that the filter paper was removed before centrifugation.

Pelleted PBMCs and fibroblasts were resuspended in 200 µL of cold methanol/water (4:1, v/v) and processed as described for cells.

Dried extracts were reconstituted in 300 µL of anhydrous acetonitrile (ACN):dichloromethane (2:1, v/v) containing 16 µmol pyridine-4-carboxylic acid, 25 µmol *N*-(3-dimethylaminopropyl)-*N’*-ethylcarbodiimide hydrochloride (EDC), and 50 µmol 4-(dimethylamino)pyridine (DMAP). Reaction mixtures were incubated at 55 °C for 4 hours with agitation at 900 rpm.

Following derivatization, 300 µL of chloroform and 150 µL of LC–MS grade water were added. Samples were vortexed at 4 °C for 10 min and centrifuged at maximum speed for 10 min at 4 °C. The lower organic phase containing the derivatized products was collected and dried in a SpeedVac. Dried samples were stored at –20 °C until analysis.

### Separation of LPG and BMP isomers by chiral HPLC column and Triple Quad MS

Dried derivatized samples were reconstituted in 50 µL of ACN, vortexed for 10 min, and centrifuged at maximum speed for 10 min at 4 °C. Finally, 45 µL of the supernatant was transferred into a glass MS vial for analysis.

2 µL of derivatized samples were analyzed using an Agilent 1290 UHPLC system coupled with an Agilent 6495D triple quadrupole platform (Agilent, Santa Clara, CA, USA). Samples were loaded and separated using a CHIRALPAK IC 83U83 column (DAICEL Chiral Technologies) held at 40 °C. Analytes were eluted by isocratic elution with 15% mobile phase A (water: methanol, 20:80, v/v, containing 10 mM ammonium acetate) and 85% mobile phase B (pure ACN) at a flow rate of 0.2 mL/min for 30 min. MS parameters were identical to those described under “Targeted lipidomics”. The precursor-to-product ion transitions (*m/z*) used for MRM are listed in **Table S3**.

Data analysis and chromatogram extraction were done using Agilent MassHunter Qualitative Analysis (v 12.0). Peak integration was done using Skyline-daily (26.1.1.209).

### Separation of GPG isomers by nanoLC-timsTOF

Dried derivatized samples were reconstituted in 50 µL ACN and then diluted with 450 µL of 0.1% formic acid in H_2_O. The samples were further desalted using a Waters Sep-Pak C18 cartridge (Waters WAT054955). The final eluate was dried down using a Speedvac and then reconstituted in 10% ACN in H_2_O with 0.1% formic acid.

2 µL of derivatized samples were analyzed using a Bruker nanoElute2 system coupled with a timsTOF HT platform (Bruker Scientific, LLC, Bremen, Germany). Chromatographic separation was performed on a C18 column (250 mm × 75 µm, 1.6 µm particle size; Ion Optics) held at 50 °C, at a flow rate of 200 nL/min. Mobile phases A and B consisted of 0.1% formic acid in water and 0.1% formic acid in ACN, respectively. The gradient increased from 5% to 95% B over 0–20 min, followed by a hold at 95% B from 20–30 min. Mass spectra were acquired in positive ESI mode over an *m/z* range of 200–1400. The inverse reduced mobility (1/*K*_0_) window had a width of 0.05 V·s/cm². Instrument settings were as follows: capillary voltage, 1500 V; dry gas flow rate, 3.0 L/min; drying temperature, 150 °C; and TIMS ramp time, 1500 ms. The accumulation time was determined by real-time ion charge control (ICC). Both the TIMS and mass calibration of the instrument were carried out using the Agilent tuning mix before sample runs. Data analysis was done using Bruker Compass DataAnalysis (v 6.2 Build 313.1.0).

### Data preparation and statistics

GraphPad Prism was used for quantitative data presentation, and figures were assembled in Adobe Illustrator 2025. All statistical analyses were performed using Prism software. Unless otherwise stated in the legends, all measurements were derived from independent samples or biological replicates. The Western blot data displayed were from representative experiments. ChemDraw was used to prepare the chemical structures depicted in **Figs. 2 and 3**.□

## Acknowledgements

We thank all members of the Abu-Remaileh laboratory for valuable discussions. We also thank Erica Forsberg, Ruben Shrestha, Liling Fang, and Robert Reed from Bruker Daltonics for their assistance and for providing access to the timsTOF instrument. We thank Lingjun Li and Shuling Xu at the University of Wisconsin-Madison for their help. We also thank Rui Zhang, Bao Li, Wen Zhang, and Liang Zhang from WuXi AppTec for their support with the custom synthesis of lipid standards. We thank Patrick Almhjell and Daniel Herschlag for helpful discussion. Additionally, we thank the Metabolomics group (directed by Dr. Yuqin Dai) of the Nucleus at Sarafan ChEM-H, Stanford University; and we thank the donors and their families for the gift of tissue that made this study possible. ChatGPT (OpenAI) and Claude (Anthropic) were used for language editing and improving manuscript clarity. All content was reviewed and approved by the authors.

## Author contributions

Conceptualization: Z.Z., J.X., M.A.-R.; Methodology: Z.Z., J.X., I.M., M.A.-R.; Investigation: Z.Z., J.X., I.M., M.J.M., A.G., C.C., A.A., M.A.-R.; Formal analysis: Z.Z., J.X., M.A.-R.; Visualization: Z.Z., J.X., I.M., M.A.-R.; Resources: X.P., N.G.-O., C.P., A.S., S.L.C., M.A.-R.; Supervision: M.A.-R.; Funding acquisition: N.G.-O., C.P., A.S., S.L.C., M.A.-R.; Writing – original draft: Z.Z., J.X., M.A.-R; Writing – review & editing: All authors.

## Data, code, and materials availability

All data is available in the main text or the supplementary materials. Plasmids and cell lines can be requested from, and requests will be fulfilled by, the corresponding author. Raw data will be deposited in a public repository upon publication.

